# Estrogen deficiency reduces maximal running capacity and affects serotonin levels differently in the hippocampus and nucleus accumbens in response to acute exercise

**DOI:** 10.1101/2024.03.05.582269

**Authors:** Earric Lee, Tuuli A. Nissinen, Laura Ylä-Outinen, Aaro Jalkanen, Jari E. Karppinen, Victoria Vieira-Potter, Arto Lipponen, Sira Karvinen

## Abstract

**Objective:** Estrogen deficiency is associated with unfavorable changes in body composition and metabolic health. While physical activity ameliorates several of the negative effects, loss of ovarian function is associated with decreased physical activity levels. It has been proposed that the changes in brain neurochemical levels and /or impaired skeletal muscle function may underlie this phenomenon.

**Methods:** We studied the effect of estrogen deficiency induced via ovariectomy (OVX) in female Wistar rats (n=64). Rats underwent either sham or OVX surgery and were allocated thereafter into four groups matched for body mass and maximal running capacity: sham/control, sham/max, OVX/control, and OVX/max, of which the max groups had maximal running test before euthanasia to induce acute response to exercise. Metabolism, spontaneous activity, and maximal running capacity were measured before (PRE) and after (POST) the surgeries. Three months following the surgery, rats were euthanized, and blood and tissue samples harvested. Proteins were analyzed from gastrocnemius muscle and retroperitoneal adipose tissue via Western blot. Brain neurochemical markers were measured from nucleus accumbens (NA) and hippocampus (HC) using ultra-high performance liquid chromatography.

**Results:** OVX had lower basal energy expenditure and higher body mass and retroperitoneal adipose tissue mass compared with sham group (p≤0.005). OVX reduced maximal running capacity by 17% (p=0.005) with no changes in muscle mass or phosphorylated form of regulatory light chain (pRLC) in gastrocnemius muscle. OVX was associated with lower serotonin metabolite 5-hydroxyindoleacetic acid (5-HIAA) level in the NA compared with sham (p=0.007). In response to acute exercise, OVX was associated with low serotonin level in the HC and high level in the NA (p≤0.024).

**Conclusions:** Our results highlight that OVX reduces maximal running capacity and affects the response of brain neurochemical levels to acute exercise in a brain region-specific manner. These results may offer mechanistic insight into why OVX reduces willingness to exercise.

## 1. Introduction

Withdrawal of ovarian hormones by ovariectomy (OVX) in rodents is commonly used to mimic the effects of menopause in the female body. Loss of ovarian function in women and female rats is associated with increased body weight and adipose tissue mass, as well as a decline in metabolic health [1–4]. However, the mediating factors are not precisely known. Due to increasing life expectancy, women can live more than one third of their lives in an estrogen deficient state [5]. Consequently, it has become more important than ever to understand the mechanisms and find solutions to counteract the possible adverse effects of menopause.

We and others have shown that physical activity ameliorates several negative effects associated with menopause in women [1,6,7]. However, the loss of ovarian hormones has been associated with decreased physical activity levels in both women and rodents [8–10]. Moreover, estrogen deficiency has also been linked to changes in brain serotonin and dopamine levels [11,12] and/or impaired skeletal muscle function [13–15]. While several studies have reported a decrease in spontaneous physical activity [11,16] and voluntary wheel running [8,17,18] after OVX in rodents, the loss of ovarian hormones and its effect on maximal running capacity is yet to be determined. Furthermore, whether alterations in muscle myosin activity or brain neurochemical marker levels are associated with potential changes in running capacity remains unexplored.

Loss of ovarian function has important effects on neurochemical marker production and release [19]. The brain dopamine system has been considered a key controller for the regulation of motivation contributing to voluntary physical activity [8,20]. Mood and motivation are associated with brain serotonin [21], which is involved in numerous physiological processes including motor and cognitive activities [22,23]. Lack of serotonin has been suggested to play a role in depression and anxiety [24], whilst acute bouts of exercise have been shown to increase brain serotonin system function [25,26]. Estrogens have been shown to increase the concentration of serotonin and to modulate serotonin action by regulating the distribution and state of its receptors [27,28]. Although an acute bout of exercise alters various neurochemical marker levels in the brain [12], whether OVX modulates this response has yet to be fully investigated. In addition, few studies have compared brain regions for how ovarian hormones and exercise affect neurochemical release, and how brain region-specific changes in neurochemicals may mediate changes in exercise behavior.

We have previously shown that loss of ovarian hormones impairs skeletal muscle function and may contribute to the loss of muscle mass associated with menopause [13,15,29]. More specifically, estrogen affects myosin regulatory light chain (RLC) activity [15,30]. According to this observation, OVX leads to a situation where fewer myosin heads are available to trigger muscle contraction. We have also shown that OVX may induce muscle atrophy through microRNA signaling, possibly leading to decreased muscle mass [29]. These changes may partly underlie the decrease in physical activity levels observed following the decline of ovarian hormones. They may also affect maximal running capacity because of reduced force production from the skeletal muscles.

Menopause is widely believed to reduce energy expenditure [31], even though recent literature challenges this topic [32,33]. At cellular level, the amount and efficiency of mitochondria and uncoupling proteins (UCP) play the main role in thermogenesis, and hence, contribute to energy expenditure [34]. UCP family contains UCP1, UCP2, and UCP3, which localize to mitochondrial inner membrane and uncouple oxidative phosphorylation leading to energy dissipation as heat [34,35]. Previous study showed that estrogen deficiency is associated with lower UCP2 mRNA expression in white adipose tissue of female rats [36]. Furthermore, it has been suggested that UCP1 is protective against metabolic dysfunction associated with loss of ovarian hormones [37].

The purpose of the present study was to investigate whether OVX reduces maximal running capacity in female rats via affecting brain neurochemical levels and/or skeletal muscle RLC activation. In agreement with previous literature, OVX led to increased body and adipose tissue masses compared with sham operation. OVX rats had similar energy intake and spontaneous activity levels, but lower energy expenditure compared with sham rats, suggesting that the adipose tissue mass increase resulted from decreased basal energy expenditure. OVX was also associated with low UCP3 level in gastrocnemius muscle. In support of our hypothesis, we report that OVX led to reduced maximal running capacity, although did not affect lower limb muscle mass or the level of the active form of RLC (pRLC) in the gastrocnemius muscle. Interestingly, we found that OVX was associated with low serotonin level in the hippocampus (HC) and high level in the nucleus accumbens (NA) in response to acute exercise, which may offer mechanistic insight into why OVX consistently reduces willingness to run in rodents.

## 2. Materials and methods

### 2.1 Ethical approval

The animal experiment was approved by the national Project Authorization Board (ELLA, Finland, permit number ESAVI/4209/2021). All procedures with the animals were conducted in accordance with the “Principles of Laboratory Animal Care’’ (NIH publication #85–23, revised in 1985) and the European Commission Directive 2010/63/EU. All efforts were made to minimize the number of animals used and their suffering.

### 2.2 Rat model

A total of 64 female Wistar (Hans) rats were purchased from Envigo (Indianapolis, USA, bred and shipped from the Netherlands). Rats arrived at the animal unit of the University of Jyväskylä at the age of 13–15 weeks (∼3 months). All rats were housed 2/cage in an environment-controlled facility (12/12 h light-dark cycle, 22^°^C) and received water and estrogen-free rodent feed (2019X, Envigo, Indianapolis, USA) *ad libitum* upon the arrival to the animal unit. During the recovery period from the surgeries (1 week) and measurements of spontaneous activity and energy and water intake, the rats were housed individually.

### 2.3 Study design

After arriving at the animal unit, the rats had one month of habituation to the new environment (Figure 1A). PRE measurements were performed when the rats were 4–6 months old. The measured outcomes were body mass, energy expenditure, fasting blood parameters (glucose, insulin), maximal running capacity, respiratory exchange ratio (RER) and spontaneous activity during 24 h metabolic cage stay (Figure 1A). Rats were then divided into sham and OVX groups matched for body mass and maximal running capacity (n=32/group). Sham and OVX surgeries were performed when the rats were 7 months old to ensure that they were fully grown before the interventions [38] (Figure 1A). Two animals were euthanized because of post-surgery complications (final n=62). After 2 months (57±4.8 days) of recovery, POST metabolism and spontaneous activity measurements were carried out prior to other follow-up measurements (Figure 1A). Thereafter, rats were further divided into control and maximal running capacity (max) groups (n=15–16/group), which were matched for body mass and maximal running capacity within a group (sham or OVX) according to the POST measurements (Figure 1B). Euthanasia and tissue harvest were performed at the age of 10–11 months, on average 3 months (104±4.8 days) after the surgery (Figure 1A). Rats in the sham group were euthanized at proestrus (determined by cytology sample) to ensure that they were in the highest possible systemic estrogen state.

**Figure 1.**
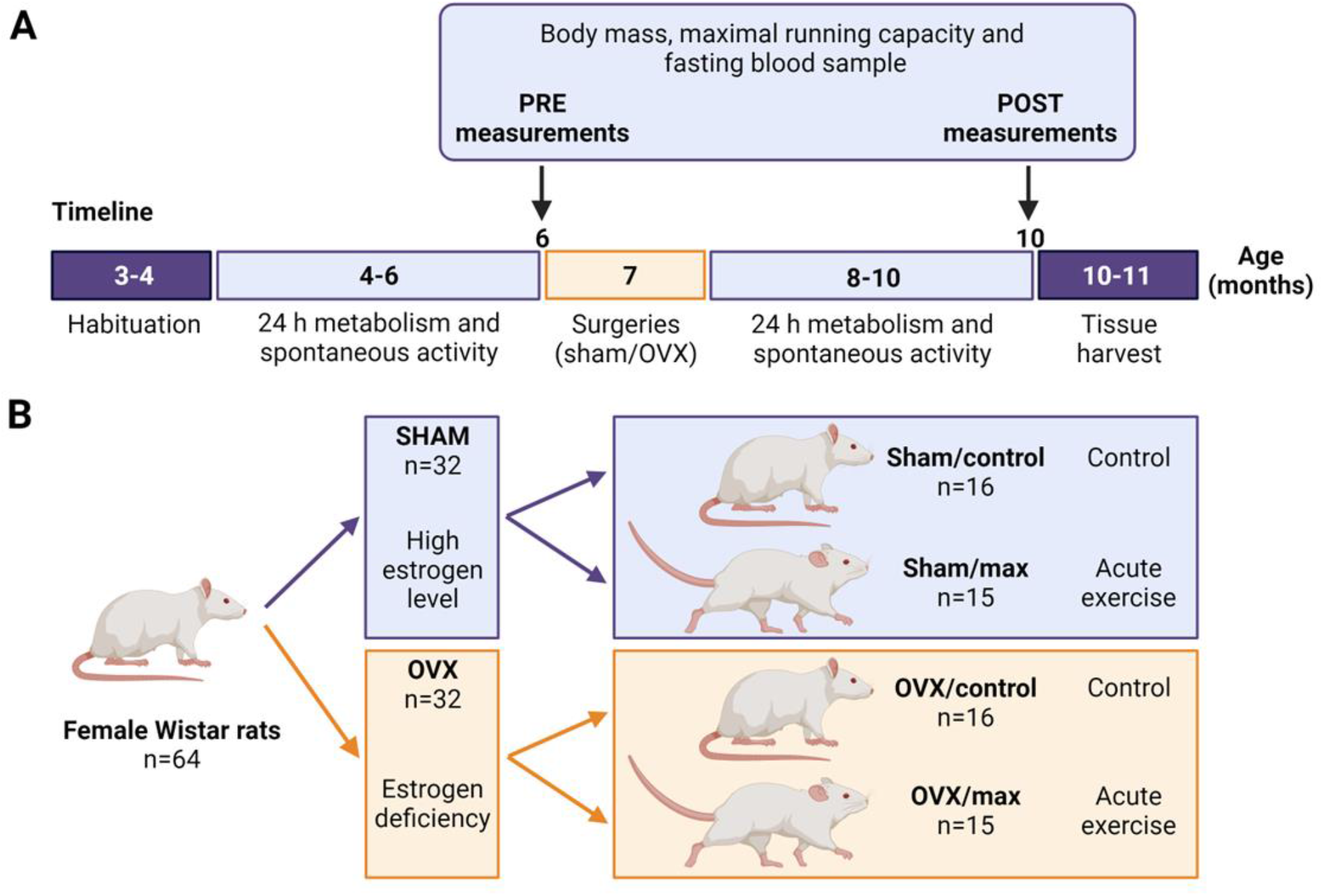
Study timeline with measurements (A) and study groups (B). Sham=ovary-intact, OVX=ovariectomy.

### 2.4 Identifying the phase of the estrous cycle

The phases of the estrous cycle (proestrus, estrus, metestrus or diestrus) were identified from vaginal mucosa samples using Giemsa stain (09204, Merck, NY, USA) and according to the appearance of nucleated or anucleated epithelial cells and leucocytes (Supplementary Figure 2B). Rats in the sham group were euthanized at proestrus to ensure high systemic estrogen levels [39,40]. The phase of the estrous cycle was also determined at the time of the PRE maximal running tests.

### 2.5 Metabolism and spontaneous activity

Energy expenditure and RER were assessed with indirect calorimetry during a 24 h metabolic cage stay as described previously [41]. Briefly, the commercial metabolic cages were equipped with Promethion^®^GA3 gas analyzers (Sable Systems, Las Vegas, NV, USA) and FR8 flow generators (Sable Systems) [41]. The flow rate was set at 3,000 ml/min and the raw data were processed using ExpeData^®^ software (Sable Systems). The Weir equation was used to calculate energy expenditure (kcal/h) = 60 × (0.003941 × V̇ O_2_ + 0.001106 × V̇ CO_2_). The metabolic cages were also equipped with ground-reaction force recording systems to record spontaneous physical activity as described previously [42]. To obtain a single value for total spontaneous activity (activity index), 1-s means were summed for the total measurement time, and the sum was divided by the body mass (kg) of the measured rat. Each rat was measured PRE and POST the sham/OVX surgery, with at least one month of recovery from the surgery before the follow-up measurement. The metabolism outcomes and spontaneous activity were determined for the whole 24 h measurement time as well as for the dark (12 h) and light (12 h) periods because rats are active during dark periods. Two OVX and one sham group rat exhibited a substantial increase in energy expenditure from PRE to POST compared with the other animals within their respective groups (>3 × interquartile range). We opted for a conservative approach and excluded these animals from the statistical analyses because the applied statistical methods were sensitive to extreme outliers.

### 2.6 Maximal running capacity and fasting blood measurements

Rats were tested for their maximal running capacity with a speed-ramped treadmill running test (15° slope, initial velocity of 10 m/min, increased 1 m/min every 2 min) at the age of 6 months as described previously [43]. Rats were first habituated to running on the treadmill with three different sessions lasting for 10 min with a low velocity (<10 m/min). Maximal running test was repeated three times with at least one day of recovery in between. The best result of the three trials (maximal running distance [m]) was considered the maximal running capacity. Maximal running capacity was assessed during PRE and POST measurements and in addition, rats allocated into max groups (sham/max and OVX/max) performed last maximal running tests immediately before euthanasia.

Since OVX rats tend to gain more adipose tissue than sham rats, the work (J) that the rats performed during the maximal running test was calculated with the following equation: Work (J) = Ascent (m) × 9.81 (N/kg) × Body mass (kg). Blood glucose was measured after 12 h fasting from blood harvested from the saphenous vein with a glucose analyzer (Hemocue® Glucose 201). Insulin was measured with ELISA from frozen (-80°C) serum samples (Mercodia, Rat Insulin ELISA). Homeostatic Model Assessment for Insulin Resistance (HOMA-IR) was calculated with the following equation: [Glucose (mmol/l) × Insulin (μg/l)]/2.43 [44].

### 2.7 Ovariectomy and sham surgeries

Surgeries (sham=ovary-intact or OVX=ovariectomy) were performed when the rats were 7 months old. Briefly, rats were first anesthetized with isoflurane with first induction with 5% in 450 ml/min of pressurized air followed by stable anesthesia with 2 % in 450 ml/min. Then, back of the animal was shaved and disinfected with repeated swapping with iodine solution (polyvinylpyrroliodine, iodine 7.5 g/kg, Jodopax, Phrmaxim AB, Helsinborg, Sweden) and 70% ethanol. One incision was done vertically at the center of the back followed by small incision at each side on fascia to reach the ovaries. When performing OVX surgery, the ovary was ligated and removed. Thereafter the fascia and skin were closed with sutures. When performing sham surgery, each step was done similarly as in OVX surgery, except that the ovaries were not removed. Rats received subcutaneous injections of buprenorphine (0.01–0.1 mg/kg, Vetergesic vet 0,3 mg/ml, Ceva Santé Animale, France) before the surgery and every 6–12 h after the surgery for the first two days and carprofen (5 mg/kg, Rimadyl vet 50 mg/ml, Zoetis Animal Health ApS, Denmark) before the surgery and every 24 h for the first 3 days after the surgery. The possible experience of pain was assessed using the Grimace scale [45]. After the surgeries, the rats were housed individually for a week to ensure that the wounds healed appropriately.

### 2.8 Body mass and energy and water intake

Body mass was followed throughout the study by weighing the rats once a week. Energy and water intakes were recorded for one week PRE and POST surgeries when the rats were housed individually. Energy and water intake were also followed between weeks 7–14 after the surgeries when the rats were housed in pairs with both rats belonging to the same study group (sham or OVX). The energy intake was calculated from the feed energy content information provided by the manufacturer (2019X, Envigo; 3.3 kcal/g). Since the rats were housed 2/cage (with a rat from the same group), energy intake was calculated as an average of two rats.

### 2.9 Neurochemical marker measurements from brain regions with UHPCL

After the animal was euthanized, the brain was immediately removed from the skull and sliced into 1 mm thick coronal brain slices with the help of the coronal brain matrix (WPI). The coronal sections were placed on cold saline-soaked filter paper on an ice bath, and brain regions were dissected using a reusable biopsy punch, 1 mm (WPI). The collected brain samples were stored in individual tubes at -20 °C until further analysis.

Seven different neurochemical markers were assayed from two brain regions, i.e., the nucleus accumbens (NA) and hippocampus (HC): serotonin, serotonin metabolite 5-hydroxyindoleacetic acid (5-HIAA), dopamine, dopamine precursor l-3,4-dihydroxyphenylalanine (L-DOPA), dopamine metabolites homovanillic acid (HVA), and 3,4-dihydroxyphenylacetic acid (DOPAC), and norepinephrine. Of the seven neurochemical markers all were detectable in the NA and four (serotonin, 5-HIAA, dopamine and norepinephrine) in the HC.

The frozen brain tissue samples were first thawed on ice and homogenized in 25 volumes (w/v) of ice-cold 0.1 M perchloric acid (Merck KGaA, Darmstadt, Germany) using an ultrasound homogenizer (Soniprep 150, MSE Scientific Instruments, Sussex, England). The homogenates were centrifuged (15000 × g, 20 min, +4 °C) and the supernatant was filtered with a 0.2 µm Acrodisc syringe filter (Pall Corporation, USA). The filtrates were further diluted with ultrapure water (NA samples 1:15 and HC samples 1:8) before analysis. The monoamine and metabolite levels were assayed using an ultra-high performance liquid chromatography (UHPLC) method with electrochemical detection (Antec Alexys Monoamine Analyzer, Antec Scientific, Hoorn, the Netherlands). The analyzer consisted of an AS 110 autosampler (set to +4 °C), two LC 110S pumps (channel 1 and channel 2), an OR 110 degasser unit with pulse damper, and a DECADE II electrochemical detector equipped with two VT-03 2 mm glassy carbon working electrodes and Ag/AgCl reference electrodes. Channel 1 was used for the analysis of dopamine and serotonin and channel 2 for the analysis of norepinephrine, L-DOPA, 5-HIAA, DOPAC, and HVA. Injection volume was 10 µL. Instrument control and data acquisition were done with Clarity data system (DataApex, Prague, the Czech Republic). The chromatographic conditions for channel 1 consisted of a C18 reversed-phase column (NeuroSep 105; 50 mm × 1.0 mm, 3 µm spherical particles, Antec Scientific) and the mobile phase (50 mmol/L phosphoric acid, 0.1 mmol/L EDTA, 600 mg/L octane sulfonic acid, 12 % v/v acetonitrile, pH was adjusted to 6.0 with 50% NaOH). Flow rate was 50 µl/min and pressure 54 bar. For separation of the analytes in channel 2, an ALF-115 column (150 mm × 1.0 mm, 3 µm, C18, Antec Scientific) was used. The mobile phase contained 50 mmol/L phosphoric acid, 50 mmol/L citric acid, 0.1 mmol/L EDTA, 600 mg/L octane sulfonic acid, pH 3.0, and 6 % v/v acetonitrile. Flow rate was 75 µl/min and pressure 194 bar. The columns and detectors were kept at 35 °C by a column oven. The applied potential was set to 0.46 V and 0.80 V for channel 1 and 2, respectively. Analyte amounts producing a detector signal with a signal-to-noise ratio >3 was considered the limit of detection (LOD). For each analyte, the linear (r^2^ >0.999) calibration curve was ranging from 0.5 to 50 nM, the lower limit of quantification (LLOQ) was 0.5 nM, and the quality control samples for low, intermediate, and high concentration levels were within the validated range of ± 15% of nominal concentrations. Before performing the statistical analysis, the extreme outliers were first excluded (>3× interquartile range).

### 2.10 Tissue and plasma harvest

At the end of the study, rats were euthanized after a 2 h fast using carbon dioxide followed by heart puncture. Tissues (heart, liver, adipose tissue, and skeletal muscles) were weighed, snap frozen in liquid nitrogen and stored at -80°C for further future analyses. Of the adipose tissue deposits, only retroperitoneal adipose tissue was harvested and weighed because of its more distinct location and uniform composition compared to visceral (i.e., omental) and/or ovarian adipose tissue deposits. Plasma (EDTA) was separated from the whole blood via centrifugation after 15 min incubation at RT (1500g, 10 min at RT) and stored as 200 µl aliquots in -80°C.

#### 2.10.1 Plasma HDL-C level

The high-density lipoprotein cholesterol (HDL-C) level was measured from the plasma of representative subsets of the groups (n=16) using an automated analyzer (Indiko Plus, Thermo Fisher, US).

#### 2.10.2. Protein analysis from skeletal muscle and adipose tissues via Western blot

Snap frozen gastrocnemius muscle and retroperitoneal adipose tissue were used for Western blot analysis as described previously (Karvinen et al., 2016). The tissue sample was dissolved in ice-cold buffer (20 mM HEPES [pH 7.4], 1 mM EDTA, 5 mM EGTA, 10 mM MgCl_2_, 100 mM, β-glycerophosphate, 1 mM Na_3_VO_4_, 2 mM DTT, 1% NP-40 [nonyl phenoxypolyethoxylethanol], 0.2% sodium deoxycholate, and 3% protease and phosphatase inhibitor cocktail [P 78443; Pierce, Rockford, IL]) and homogenized using a stainless steel bead in TissueLyser II (Qiagen, Germany) for 2 min at 30 Hz. Samples were mixed for 30 min in end-over-end rotation at +4 °C. The tissue homogenate was then centrifuged at 10,000 x g for 10 min at +4°C. Total protein content was determined using the bicinchoninic acid protein assay (Pierce Biotechnology, Rockford, IL) with an automated KoneLab instrument (Thermo Scientific, Vantaa, Finland).

Aliquots of tissue homogenate were solubilized in Laemmli sample buffer and heated at 95°C to denaturate proteins, except for detecting mitochondrial protein complexes vis Total OXPHOS Cocktail, when samples were heated at 37°C. Samples containing 30 μg of total protein were separated by SDS-PAGE for 30 min at 270 V using 4–20% gradient gels on Criterion electrophoresis cell (Bio-Rad Laboratories, Richmond, CA). Proteins were transferred to nitrocellulose membranes, which were blocked in blocking buffer (Licor, Cat no 927-70001) for 2 h and then incubated overnight at +4°C with primary antibodies in 1:1 TBS and blocking buffer to analyze the content of pRLC (1:1000; ab2480, Abcam), Total OXPHOS Cocktail (OXPHOS, 1:1000; ab110413; Abcam), UCP1 (1:1000; ab10983, Abcam), UCP2 (1:500; ab67241 Abcam), and UCP3 (1:1000; ab3477 Abcam). The proteins pRLC, OXPHOS, UCP2, and UCP3 were measured from gastrocnemius muscle and the proteins OXPHOS, UCP1, and UCP2 were measured from retroperitoneal adipose tissue.

After the primary antibody incubation, membranes were washed in TBS-Tween (TBS-T, 0.1%), and then incubated with a suitable secondary antibody diluted in TBS-T with 1:1 blocking buffer for 1 h followed by washing in TBS-T. Proteins were visualized using fluorescent secondary antibodies and quantified using ChemiDoc MP device (Bio-Rad Laboratories, Hercules, CA, USA) and quantified with Image Lab software (version 6.0; Bio-Rad Laboratories, Hercules, CA, USA). First, the target and total protein and levels were normalized to the membrane average to eliminate differences in signal intensity between the membranes. Thereafter, target protein levels were normalized to corresponding total protein amount (full lane) and results are shown in arbitrary units (AU).

### 2.11 Statistics

The summary statistics are presented as group means with standard deviations. The normality of variables was assessed using the Shapiro-Wilk test followed by Levene’s test for examining the equality of the variances. When the normality criteria were met, the differences between groups were examined using Student’s *t*-test and the difference between PRE and POST measurements using Student’s paired *t*-test. OVX and sham groups exhibited different variances in RER and thus, for this variable, comparisons were performed using Welch’s *t*-test. When the normality criteria were not fulfilled, the differences between groups were examined using Mann-Whitney U-test and the difference between PRE and POST measurements using Wilcoxon signed-rank test. The main effects of OVX, acute exercise and their interaction (OVX × Acute exercise) were determined using two-way analysis of variance (ANOVA). Energy expenditure is highly dependent on body mass and regression-based methods are needed particularly when comparing groups with differing masses [47]. Therefore, changes in energy expenditure outcomes between groups were also compared using linear mixed-effect models (*nlme* package) with animal identification as the random effect, group × time interaction as the explanatory variable, and body mass as a covariate. For the analysis of changes in 24 h energy expenditure, a separate model also included energy intake and spontaneous activity as covariates. Correlation analyses were performed and visualized using *metan* package. In all analyses, a p-value ≤0.05 was considered to indicate statistical significance. Linear mixed-effect models and correlation analysis were generated using R version 4.3.1.

## 3. Results

### 3.1 OVX led to increased body mass and decreased maximal running capacity

Before the surgeries, the rats were divided into sham and OVX groups matched for body mass and maximal running capacity (n=32/group). Accordingly, the sham and OVX groups did not differ in their body mass or maximal running capacity at PRE measurements (sham vs. OVX p≥0.642, Figure 2A-B). In line with previous literature, OVX rats were heavier than sham rats at POST measurements (sham vs. OVX, p≤0.01, Figure 2A). On average, sham group gained weight by 12% vs. 25% in the OVX group throughout the study (p≤0.001, Figure 2A). The weekly body mass tracking revealed a significant difference in body mass between sham and OVX groups starting three weeks post-surgery (Supplementary Figure 1A).

**Figure 2.**
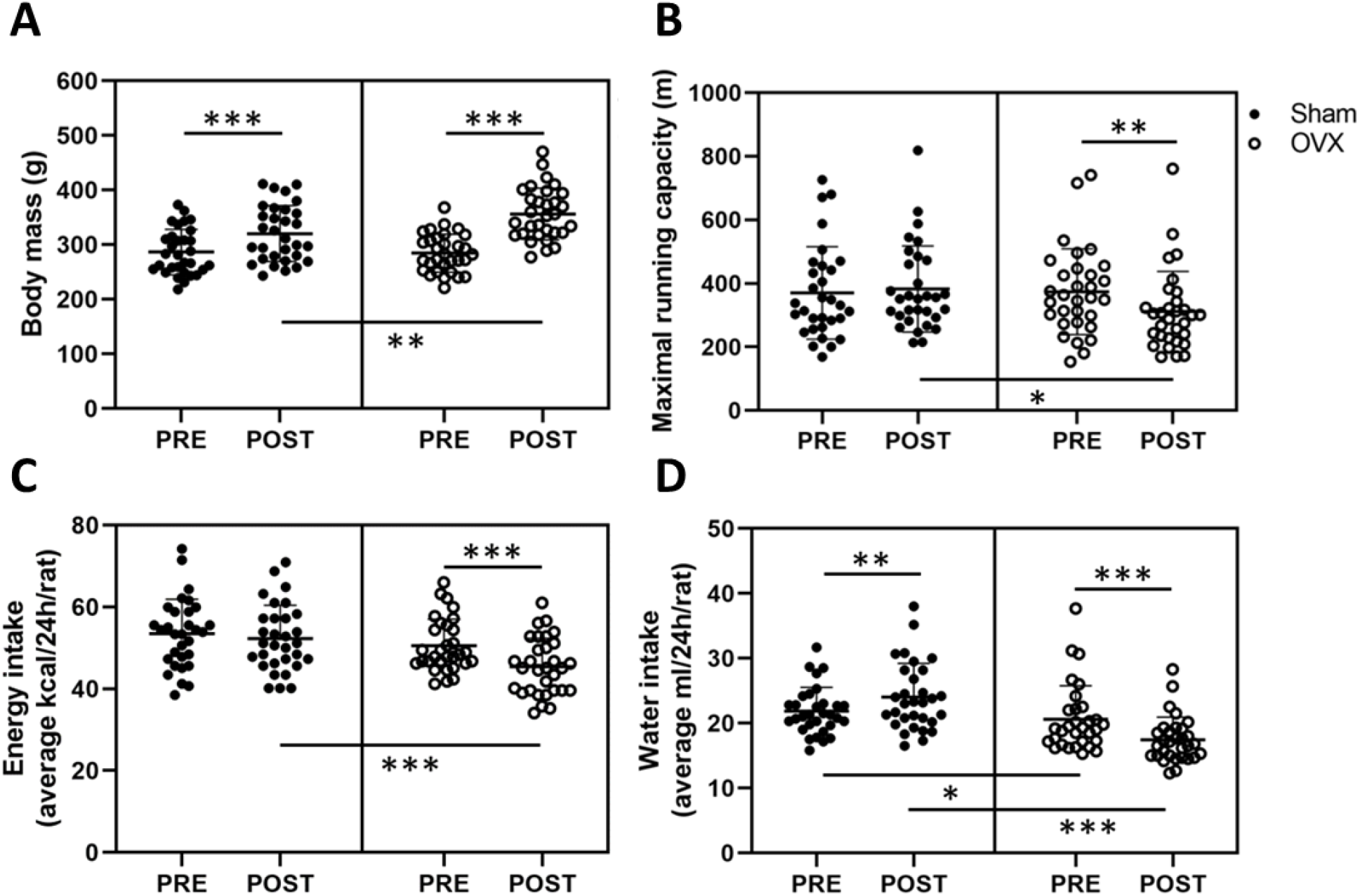
Body mass (A), maximal running capacity (B), energy intake (C), and water intake (D) before (PRE) and after (POST) the sham/OVX surgeries. Figures show individual samples with mean and SD. *p≤0.050, **p≤0.010, and ***p≤0.001.

Prior to the surgeries, the estrous cycle phase distribution in the whole study population during the best running trial was the following: proestrus 23%, estrus 23%, metestrus 16%, and diestrus 39%, indicating that maximal running capacity was not associated with higher estrogen level (i.e. proestrus). At POST measurements, OVX group had lower maximal running capacity than the sham group (sham vs. OVX p=0.013, Figure 2B). On average OVX rats reduced their maximal running capacity by ∼17% (p=0.005, Figure 2B). There were no differences between the sham and OVX groups in work (J) performed during the maximal running tests in PRE or POST measurements [PRE sham: 300 (109), OVX: 295 (83); POST sham: 341 (107), OVX: 310 (95), p≥0.245], yet within sham group the increase in work at POST vs. PRE measurements was significant (p=0.008).

The sham and OVX groups did not differ in average daily energy intake (p=0.174, Figure 2C) at PRE measurements, whereas at POST measurements, the average energy intake of OVX group had decreased (p<0.001, Figure 2C). Rats in the sham group had higher water intake than rats in the OVX group both PRE and POST surgeries (p≤0.025). Water intake post-surgery increased in the sham group and decreased in the OVX group (p≤0.004, Figure 2D). The energy intake did not differ between sham and OVX groups in weeks 7–13 (p≥0.139) while the OVX group had lower energy intake at 14 weeks (p=0.011, Supplementary Figure 1B). Consistent with the water intake measured from individual rats, rats in the OVX group had lower water intake between weeks 7–14 (p≤0.031, Supplementary Figure 1C).

### 3.2 OVX was associated with higher fat mass and higher HDL-C level

In addition to having higher body mass, the OVX group had higher retroperitoneal adipose tissue mass and lower liver mass compared with the sham group (p≤0.005, Table 1). The uterine mass of rats in the sham group was larger compared with the OVX group, which affirms the success of the OVX surgeries (p≤0.001, Table 1). Picture of representative uteri at the end of the study are shown in Supplementary Figure 2A. There were no differences in the individual muscle masses between the groups, yet OVX group had lower total muscle mass/body mass ratio compared with the sham group (p≤0.001, Table 1).

**Table 1.**
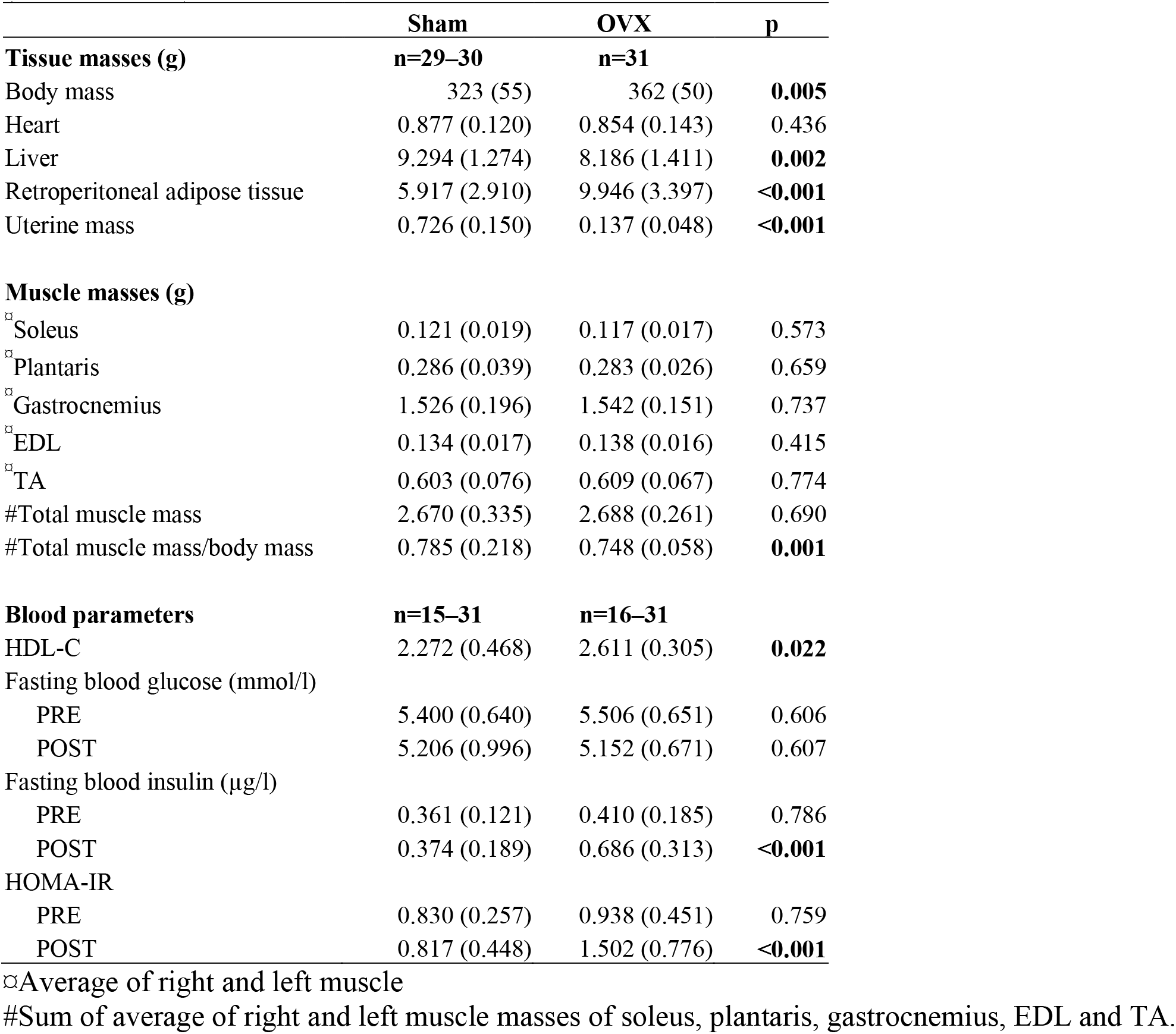
Body and tissue masses at the end of the study and blood parameters of sham and OVX rats (mean with SD).

The blood parameters revealed that OVX rats had higher plasma HDL-C level compared with sham rats (p=0.022, Table 1). The fasting blood glucose did not differ between the sham and OVX groups PRE or POST surgeries (p≥0.606, Table 1), yet OVX group had higher fasting insulin and HOMA-IR levels at POST measurements compared with sham group (p<0.001).

### 3.3 OVX led to decreased energy expenditure

At PRE measurements, rats allocated to the sham and OVX groups did not differ in energy expenditure, RER, spontaneous activity level, and energy or water intake (p≥0.129, Table 2). Even though the OVX rats were heavier at POST measurements, their absolute energy expenditure was lower, particularly during dark hours (Table 2). When adjusted for body mass, 24 h energy expenditure decreased by -0.22 kcal/h (95% CI -0.30 to -0.15, p<0.001) in the OVX group compared with the sham group. When examining the light and dark hours separately, energy expenditure during light hours decreased by -0.15 kcal/h (95% CI -0.22 to -0.08, p<0.001) and during dark hours by -0.30 kcal/h (95% CI -0.40 to -0.20, p<0.001) in the OVX group compared with the sham group. Furthermore, when adjusted for body mass, energy intake and spontaneous activity, 24 h energy expenditure decreased by -0.20 kcal/h (95% CI -0.28 to -0.12, p<0.001) in the OVX group compared with the sham group. RER and spontaneous activity decreased in both groups during the study and were similar between groups at PRE and POST measurements (Table 2). Both energy and water intake decreased in the OVX group (p<0.001, Table 2) and at POST measurements, the OVX group consumed less water than the sham group (p<0.001, Table 2).

**Table 2.** Energy expenditure, RER, spontaneous activity, and energy and water intake of sham and OVX rats (mean with SD).

|  | PRE |  |  |  | POST |  |  |  | Change |  |  |  |
| --- | --- | --- | --- | --- | --- | --- | --- | --- | --- | --- | --- | --- |
|  | SHAM | OVX | Difference | p-value | SHAM | OVX | Difference | p-value | SHAM | p-value | OVX | p-value |
| **Energy expenditure (kcal/h)** |  |  |  |  |  |  |  |  |  |  |  |  |
| 24 h | 1.77 (0.19) | 1.75 (0.19) | 0.020 | 0.728 | 1.92 (0.17) | 1.77 (0.16) | 0.150 | < 0.001 | 0.16 | < 0.001 | 0.02 | 0.331 |
| Light h | 1.46 (0.14) | 1.46 (0.16) | 0.000 | 0.982 | 1.66 (0.13) | 1.58 (0.15) | 0.070 | 0.053 | 0.19 | < 0.001 | 0.12 | < 0.001 |
| Dark h | 2.07 (0.26) | 2.03 (0.24) | 0.040 | 0.580 | 2.19 (0.22) | 1.96 (0.19) | 0.230 | < 0.001 | 0.12 | < 0.001 | -0.08 | 0.018 |
| **Respiratory exchange ratio (RER)** |  |  |  |  |  |  |  |  |  |  |  |  |
| 24 h | 0.87 (0.06) | 0.89 (0.03) | 0.020 | 0.168 | 0.83 (0.06) | 0.84 (0.03) | 0.000 | 0.884 | -0.04 | 0.003 | -0.05 | < 0.001 |
| Light h | 0.84 (0.06) | 0.85 (0.04) | -0.010 | 0.300 | 0.81 (0.06) | 0.81 (0.03) | 0.000 | 0.824 | -0.03 | 0.019 | -0.04 | < 0.001 |
| Dark h | 0.90 (0.06) | 0.93 (0.04) | -0.020 | 0.137 | 0.86 (0.07) | 0.86 (0.03) | 0.000 | 0.949 | -0.05 | 0.001 | -0.07 | < 0.001 |
| **Spontaneous Activity (AU)** |  |  |  |  |  |  |  |  |  |  |  |  |
| 24 h | 0.057 (0.020) | 0.053 (0.019) | 0.003 | 0.459 | 0.042 (0.015) | 0.045 (0.020) | -0.003 | 0.522 | -0.015 | 0.003 | -0.008 | 0.029 |
| Light h | 0.027 (0.012) | 0.026 (0.011) | 0.002 | 0.456 | 0.021 (0.010) | 0.025 (0.012) | -0.003 | 0.247 | -0.005 | 0.072 | 0 | 0.970 |
| Dark h | 0.086 (0.032) | 0.081 (0.028) | 0.006 | 0.496 | 0.062 (0.021) | 0.065 (0.030) | -0.003 | 0.688 | -0.024 | 0.001 | -0.016 | 0.004 |
| **Energy intake (kcal)** |  |  |  |  |  |  |  |  |  |  |  |  |
| 24 h | 42.0 (13.2) | 45.0 (11.9) | -3.0 | 0.361 | 41.1 (14.9) | 36.5 (8.6) | 4.6 | 0.150 | -0.8 | 0.790 | -8.5 | < 0.001 |
| **Water intake (ml)** |  |  |  |  |  |  |  |  |  |  |  |  |
| 24 h | 21.9 (7.0) | 19.3 (6.0) | 2.6 | 0.129 | 23.1 (6.4) | 14.1 (4.7) | 9.1 | < 0.001 | 1.3 | 0.362 | -5.2 | < 0.001 |

### 3.4 Acute exercise was associated with low serotonin level in the hippocampus (HC) and high level in the nucleus accumbens (NA) only among OVX rats

Before the euthanasia, the rats in the sham and OVX groups were divided into control and maximal running test (max) subgroups matched for body mass and maximal running capacity (n=15– 16/group) (Figure 1B). Consistent with this, the groups (sham/control vs. sham/max and OVX/control vs. OVX/max) did not differ in their body mass or maximal running capacity (p≥0.734, Supplementary Figure 3A-B), yet body mass was higher and maximal running distance lower in the OVX compared with sham at POST measurements (p≤0.013, Supplementary Figure 3A-B). The running distance the rats performed right before euthanasia did not differ between the groups (sham/max vs. OVX/max, p=0.299, Supplementary Figure 3C).

All seven measured neurochemical markers were detected in the NA (Figures 3 A-B, Supplementary Figures 4A-E). OVX was associated with lower serotonin metabolite, 5-HIAA, level in NA compared to sham (p≤0.007, Figure 3E). Interestingly, acute exercise was associated with an increase in NA serotonin level in OVX rats (p=0.036), but not in sham rats (Figure 3A, E).

**Figure 3.**
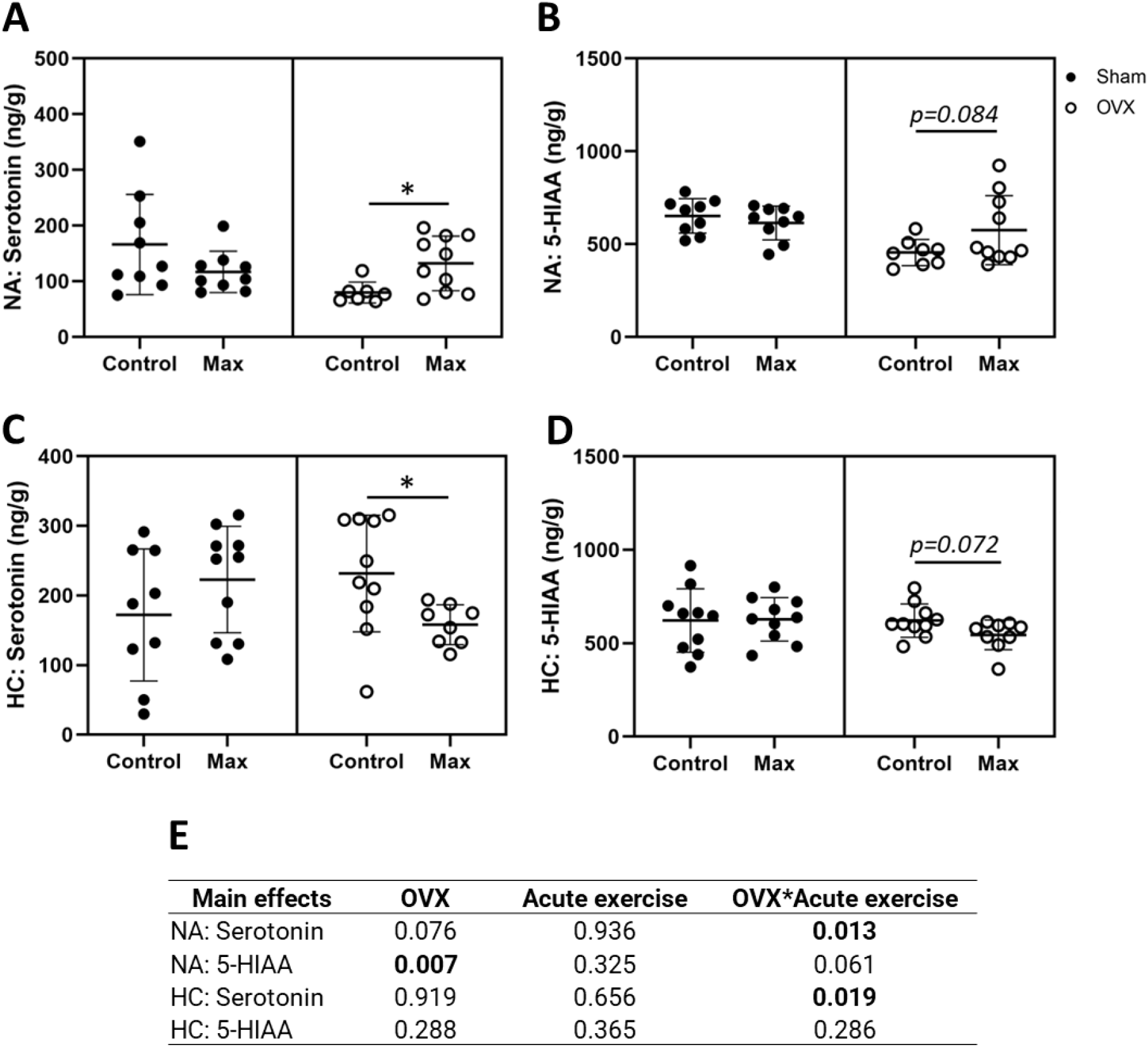
Serotonin (A) and serotonin metabolite 5-HIAA (B) in the nucleus accumbens (NA) and in the hippocampus (HC) (C,D). The main effects of OVX, acute exercise and their interaction on the measured neurochemical markers (E). Figures show individual samples with mean and SD. *p≤0.050.

Of the seven measured neurochemical markers, four were detected in the HC: dopamine, serotonin, norepinephrine, and the serotonin metabolite 5-HIAA (Figures 3 C-D, Supplementary Figures 4E-F). In contrast to NA, neither serotonin nor 5-HIAA levels were significantly affected by OVX in HC (Figure 3C, D). However, unlike the case with NA, acute exercise was associated with a reduction in HC serotonin level (p=0.024, Figure 3C) in OVX animals only. Uncertainty remains whether acute exercise is also associated with elevated 5-HIAA levels in OVX animals (p=0.084, Figure 3B).

We also examined the effects of OVX, acute exercise, and their interaction on serotonin/dopamine and serotonin/norepinephrine ratios but observed no significant effects (p≥0.056, Supplementary Figure 5). The closest to a significant observation was the interaction of OVX and acute exercise in NA serotonin/dopamine ratio (p=0.056, Supplementary Figure 5A).

Together with the significant interaction of OVX and acute exercise on serotonin in both NA and HC our results indicate that OVX rats have a distinct response to exercise in these two brain regions.

### 3.5 OVX was associated with lower UCP3 level in gastrocnemius muscle

There were no effects of OVX, acute exercise, or their interaction on pRLC level in the gastrocnemius muscle (p≥0.153, Figures 4A, E). However, OVX was associated with lower UCP3 level (p=0.022), which was not affected by exercise in either group. There was an interaction of OVX and acute running for UCP2 level in gastrocnemius level (p=0.018) such that exercise reduced UCP2 in OVX rats only (p=0.023, Figures 4C-E).

**Figure 4.**
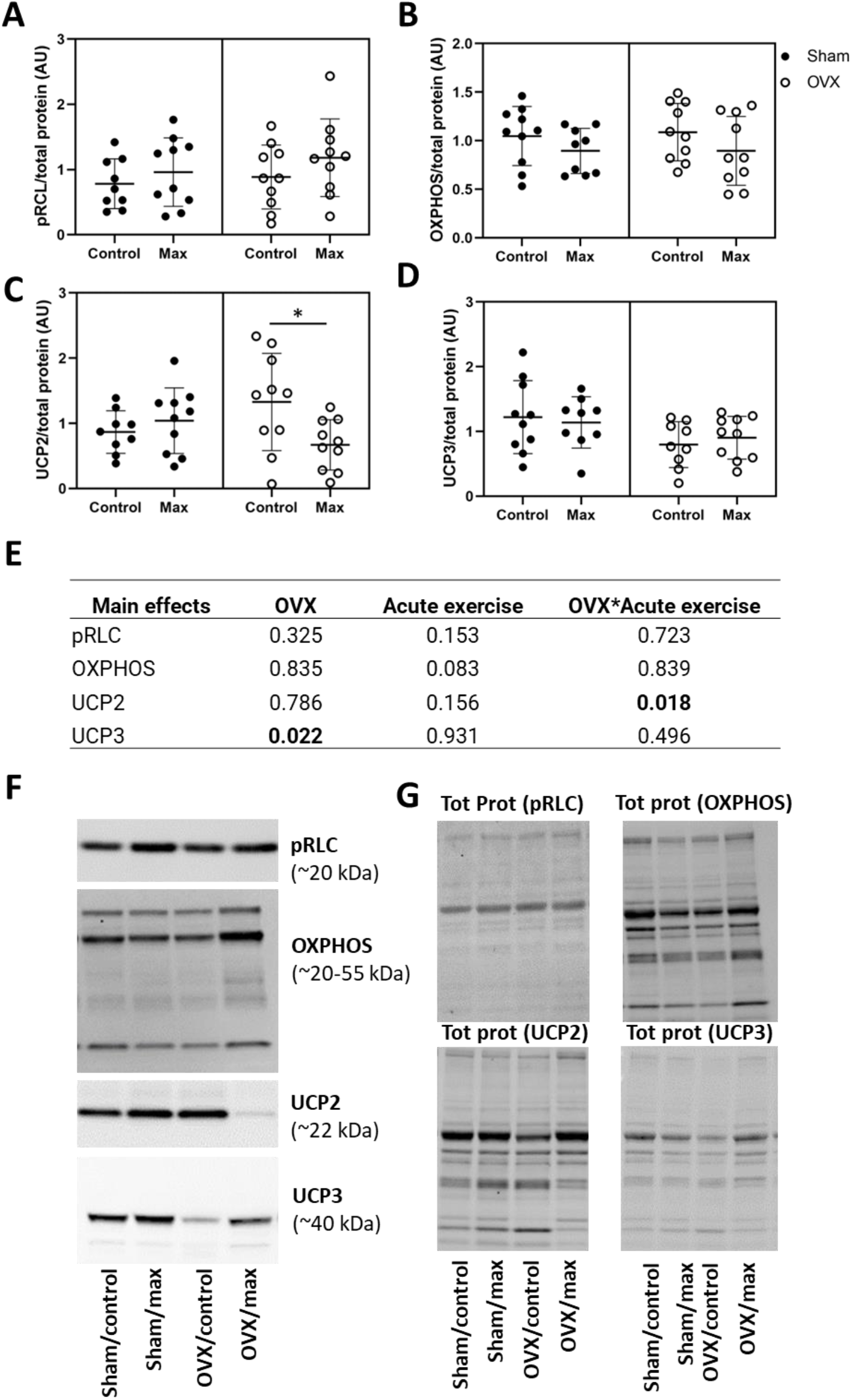
Protein levels of pRLC (A), OXPHOS (B), UCP2 (C), and UCP3 (D) relative to total protein level in gastrocnemius muscle with representative blot images (F) and corresponding blot images of total protein content (G). The main effects of OVX, acute exercise and their interaction on the measured proteins (E). Tot prot=total protein. Figure shows individual samples with mean and SD. *p<0.050.

### 3.6 OVX did not affect the level of studied proteins in retroperitoneal adipose tissue

We observed no effects of OVX, acute exercise or their interaction in the protein levels of OXPHOS, UCP1 or UCP2 in retroperitoneal adipose tissue (p=0.060, Figure 5).

**Figure 5.**
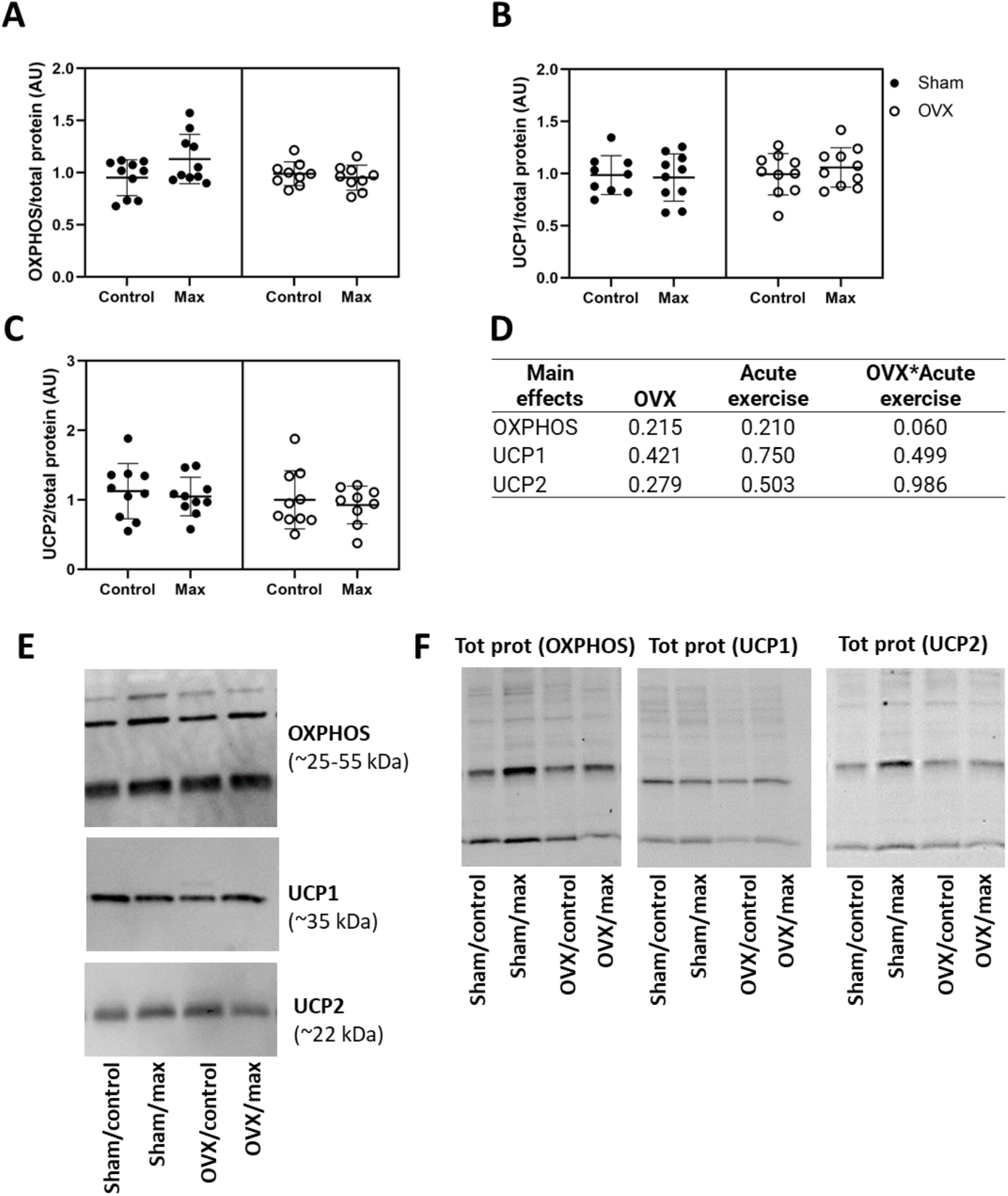
Protein levels of OXPHOS (A), UCP1 (B), and UCP2 (C) relative to total protein level in retroperitoneal adipose tissue with representative blot images (E) and corresponding blot images of total protein content (F). The main effects of OVX, acute exercise and their interaction on the measured proteins (D). Figure shows individual samples with mean and SD. Tot prot=total protein.

### 3.7 Maximal running capacity was positively associated with serotonin and 5-HIAA levels in the nucleus accumbens (NA)

To gain additional insight into the physiological relevance of OVX’s effects on brain region-specific responses to exercise, correlations among neurochemical markers and parameters of health, performance and metabolism were assessed.

When examining the correlations within the whole study population, we observed a positive correlation between maximal running capacity (POST) and dopamine level in the HC, and serotonin, DOPAC, 5-HIAA, and HVA levels in the NA (p≤0.048, Figure 6). Also, muscle pRLC protein level had a positive correlation with maximal running capacity examined immediately prior to euthanasia (END, p=0.017). Norepinephrine level in the NA was negatively correlated with body mass and retroperitoneal adipose tissue mass (p≤0.009, Figure 6), while 24 h energy expenditure (kcal/h) had a positive correlation with dopamine level in HC and 5-HIAA level in NA as well as OXPHOS and UCP2 protein levels measured from the retroperitoneal adipose tissue (p≤0.050, Figure 6). Interestingly, L-DOPA level in the NA had a positive correlation with retroperitoneal fat mass (p=0.044), but not with body mass (p=0.124, Figure 6). In addition, adipose tissue OXPHOS protein level had positive correlations with dopamine, serotonin and norepinephrine levels in the HC (p≤0.037, Figure 6). Expectedly, body mass and retroperitoneal fat mass had a positive correlation (p<0.001). Additionally, maximal running capacity was negatively correlated with body mass and retroperitoneal fat mass (p≤0.002, Figure 6). We observed no significant correlation between the maximal running capacity results (POST, END) and serotonin/dopamine ratios in NA or HC (p≥0.160, Figure 6). Serotonin and its metabolite 5-HIAA had a positive correlation both in NA and HC (p≤0.005, Figure 6), indicating robust measurement of the brain neurochemical markers.

**Figure 6.**
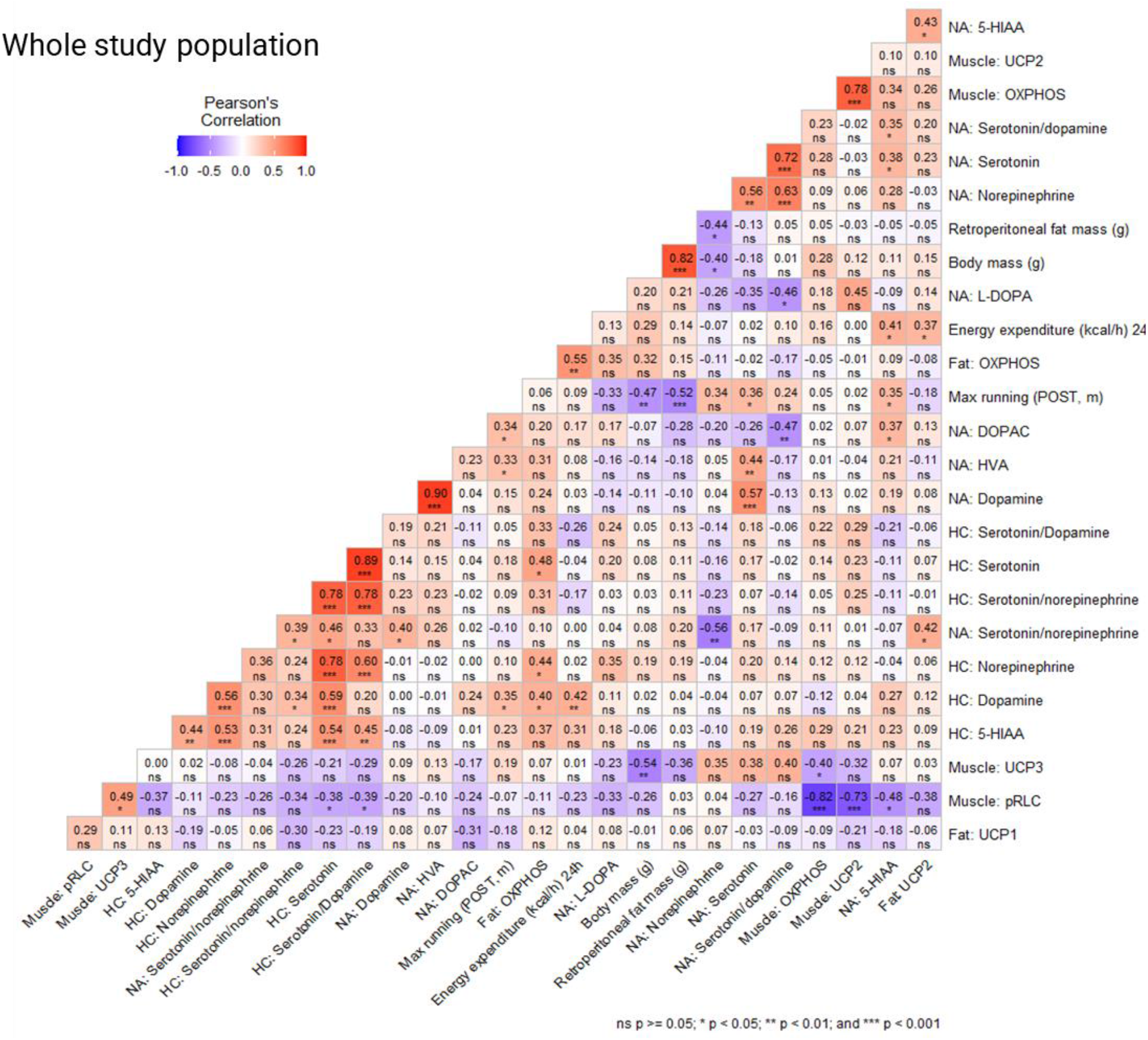
Correlation coefficients (Pearson’s correlation) in the whole study population of brain neurochemical markers in the nucleus accumbens (NA) and hippocampus (HC), protein levels in muscle (gastrocnemius), adipose tissue (retroperitoneal deposit), body mass, retroperitoneal fat mass, maximal running capacity (POST, END), and 24h energy expenditure (kcal/h).

We examined the same correlations in the following study populations separately: sham, OVX, control (no acute exercise) and max (acute exercise) and observed minor differences between the study populations (Supplementary Figures 6-9). In the sham group, body mass had a positive correlation with OXPHOS protein level measured from gastrocnemius muscle (p≤0.016, Supplementary Figure 6), while in the OVX group body mass had a positive correlation with OXPHOS and UCP2 protein levels measured from retroperitoneal adipose tissue (p≤0.032, Supplementary Figure 7). Interestingly, pRLC level in the muscle was positively correlated with maximal running capacity (END) in the sham group, but not in OVX group (p=0.036 and p=0.145, respectively, Supplementary Figure 6). OVX group had a negative correlation between maximal running capacity (END) and serotonin/dopamine ratio in HC (p=0.011, Supplementary Figure 7). In the control group there was a positive correlation between maximal running capacity (POST) and serotonin/dopamine ratio in NA (p=0.036, Supplementary Figure 8). In the max (acute exercise) group the maximal running capacity (END) had a negative correlation with L-DOPA level in NA (p=0.021) and a strong positive correlation with max running capacity (POST). This suggests that the maximal running tests performed immediately before euthanasia had similar results with maximal running tests performed at time point POST (p=0.001, Supplementary Figure 9).

## 4. Discussion

In this study, age-appropriate adult female rats were ovariectomized (OVX) or underwent sham operations to assess the effect of age-related ovarian hormone loss (i.e. mimicking menopause in women) on metabolism and body composition, as well as skeletal muscle, adipose tissue, and brain region-specific responses to an acute exercise stimulus. We show that in addition to causing weight gain due to reduced energy expenditure, OVX led to reduced maximal running capacity despite no changes in lower limb muscle mass or the pRLC level in the gastrocnemius muscle. Above all, we report for the first time that OVX affected the brain’s response to exercise in a brain region-specific manner. These findings may offer novel mechanistic insight into the role of specific brain regions in mediating OVX-induced changes in ones’ willingness to run.

### 4.1. OVX increased adiposity and reduced energy expenditure

In line with previous literature, OVX led to increased adiposity, and when measured ∼8 weeks post-surgeries, a reduced total energy expenditure. Unsurprisingly, OVX rats were heavier than sham-operated rats [3,4,48]. However, this occurred in the absence of significant differences in spontaneous physical activity, which appears contradictory to other published works [17,18,49].In the present study, the difference in body mass between the groups became apparent by three weeks post-surgeries. Plausible explanations are increased energy intake, decreased spontaneous activity (i.e., activity-related energy expenditure) and/or decreased basal energy expenditure.

Our data support reduced energy expenditure, and not increased energy intake or decreased spontaneous activity, as the main cause of adiposity increase. While the sham and OVX groups did not differ in daily energy intake prior to surgeries, energy intake of the OVX group declined after surgery despite weight gain. Based on the current evidence it remains unclear whether OVX-induced weight gain is due to reduced energy expenditure or increased energy intake. Some previous studies have observed an increase in energy intake following OVX in rats [3,4,48]. However, in these studies there was only a short-term follow-up (∼1-month post-surgery) on energy intake. Hence, it is possible that the energy intake ultimately declined in those studies as well. Indeed, McElroy and Wade (1987) reported that OVX increased energy intake in a transient manner such that intake increased during the first five weeks after the surgery but normalized to the level of controls during weeks 6–10 [50]. Importantly, despite the similar energy intake, the body weights of the OVX rats remained 12–16% above sham group, similar to what we observed in the current study.

Regarding energy expenditure, we observed no significant differences in spontaneous activity levels between the groups during the 24 h metabolic measurements before or after the surgeries. This is contrary to several other studies that have reported substantial decreases in voluntary wheel running activity in rats after OVX [17,18,49]. A decrease in voluntary spontaneous activity in OVX rats compared with sham rats both immediately [16] and 6 weeks after the surgeries [51] have been reported. The 24 h assessment period used in the present study may have been too short to detect differences, since some studies use longer (e.g., 72-h or greater) assessment period. We did, however, detect differences in energy expenditure, which decreased in the OVX group. This was present even after controlling for body mass, energy intake and spontaneous activity, providing robust evidence that the loss of ovarian hormone stimulation decreased basal energy expenditure. An earlier study observed an increase in energy intake during the early period of OVX-induced weight gain (within 3 weeks of surgery), while only a minor change in total energy expenditure was detected [16]. Studies in mice also support that OVX decreases energy expenditure without altering energy intake [52]. It is thus likely that OVX-mediated reduction in energy expenditure occurs more slowly and may not be detected until several weeks following surgery.

Mechanistically, decreased fat oxidation has been proposed to explain menopause-associated weight gain [53]. However, we found no differences in respiratory exchange ratio (RER) values, as an index of relative fat oxidation, between the sham and OVX groups. This finding is expected because the animals were in energy balance and were fed same chow. Based on laws of conservation of energy and mass, RER between groups will be identical in this case [54]. OVX did not alter substrate partitioning in the present study, and the reduction in energy expenditure might not be explained by decreases in fat oxidation. It is also possible that our system was not sensitive enough to detect subtle differences in RER over the 24 h time period of assessment. Regardless, our results suggest a decrease in basal energy expenditure may contribute to the increased body mass caused by OVX. The exact mechanisms are somewhat vague, and we are also uncertain of how well these data translate to humans, since whether menopause directly leads to reduced basal energy expenditure in middle-aged women remains unknown [33].

Regarding mechanisms by which loss of ovarian hormones in rodents or women results in changes in energy expenditure, there are a growing number of studies suggesting that estrogen may affect uncoupling proteins (UCP), which dissipate energy at the cellular level by uncoupling oxygen utilization from ATP production [34]. Indeed, estrogen-sufficient rodents and humans have been shown to have greater UCP1 expression in adipose tissue compared to age-matched males [55–57]. Moreover, a reduction in UCP1 expression is associated with a reduction in basal metabolism [37]. In the present study, the rats were housed below thermoneutrality, which likely affects the level of UCPs; we measured UCPs 1, 2, and 3 in skeletal muscle and adipose tissue.

Compared to the sham group housed under the same temperature conditions, OVX did not appear to significantly affect UCP1 or UCP2 yet was associated with lower UCP3 protein content in skeletal muscle. UCP3 is the main uncoupling protein expressed in skeletal muscle and is known to play a significant role in non-shivering thermogenesis [34,58]. Moreover, mice overexpressing UCP3 in skeletal muscle weigh less, have a decreased amount of adipose tissue, are protected from fat-induced insulin resistance, and have an increased resting oxygen consumption [59,60]. Thus, the lower expression of UCP3 in OVX rats may have contributed to their increased fat mass and decreased energy expenditure. Indeed, skeletal muscle UCP3 was negatively associated with body mass. Although not significantly affected by OVX, the adipose tissue levels of UCP2 and mitochondrial OXPHOS proteins were positively associated with energy expenditure. This supports the growing body of literature suggesting that white adipose tissue contributes significantly to resting energy expenditure [61,62], especially among females who have greater relative adiposity. While these data are intriguing and warrant further study, it is likely that a variety of factors (e.g., rodent line/strain, age at OVX, type of diet, etc.) influence the changes in energy balance and energy expenditure that occur with loss of ovarian hormones. Future studies should carefully control for such factors to obtain a better understanding of their interplay.

### 4.2. OVX increased adiposity and caused metabolic dysfunction

In addition to having higher body mass, the OVX group had higher retroperitoneal adipose tissue mass (i.e., a visceral depot) and lower liver mass compared with the sham group, in agreement with previous studies [4,63]. The increase in body fat also concurs with findings from human studies done by our group [1] and others [53,64], demonstrating an accelerated increase in adiposity during menopause. Estrogen deficiency with a concurrent increase in adipose tissue may lead to a greater risk of metabolic diseases, such as type 2 diabetes [65,66].

Some studies have observed a significant increase in fasting blood glucose after OVX [67,68], conforming with the observation that estrogen deficiency results in declined insulin-stimulated glucose disposal [69]. We did not observe a difference in fasting glucose levels, but the OVX group had higher fasting insulin levels as well as higher HOMA-IR than the sham group, indicating impaired insulin action. This observation corresponds to previous findings showing exacerbation of insulin resistance with estrogen deficiency [67,69].

In the present study, the OVX group had higher HDL-C compared with the sham group, which has been observed before [70]. It has been suggested that the atheroprotective effect of HDL may be weaker in women after menopause [71,72]. Previously we [6] and others [72,73] have reported a higher HDL-C level following menopause in women [71,72]. This previously observed increase in HDL-C level following menopause is replicated in our present animal study and thus validating prior findings.

### 4.3. OVX reduced maximal running capacity

Expectedly, OVX reduced maximal running capacity. Firstly, we ruled out an effect of current circulating estrogen level on running by examining the effect of estrous cycle phase on the best running trial. Similar to previously published data in women and rodents [74,75] observed no association between the highest estrogen phase (i.e. proestrus) and best running trial among sham rats. However, as predicted, long-term estrogen deficiency caused by OVX led to lower maximal running capacity compared with the sham treatment. We then examined two possible contributors to the observed decrease in running capacity: phosphorylation of regulatory light chain of myosin (pRLC) protein content in skeletal muscle (i.e., a physiological change), and brain neurochemical markers (i.e., a neurochemical change).

It has previously been shown that estradiol modulates RLC activity in skeletal muscle (EDL) in mice [76]. Specifically, OVX-operated C57BL/6 female mice had lower pRLC level in EDL muscle compared with sham-operated mice [76]. However, we observed no differences between sham and OVX groups in the pRLC level in gastrocnemius muscle. Mouse EDL muscle is comprised of fast type (type II) muscle fibers (<1% type I fibers) whereas rat gastrocnemius muscle comprises of both fast and slow type (type I) muscle fibers (∼10% type I fibers) [77,78]. Thus, the difference in the muscle fiber type studied may partly explain the observed difference in the results. However, with regards to running, the gastrocnemius muscle is the largest lower limb muscle contributing to locomotion. Therefore, RLC phosphorylation might not be the key contributor to the observed decline in maximal running performance in the present study. We did find that pRLC protein level was positively associated with maximal running capacity (END), yet this observation was true only for the sham group. These results are in support of exercise stimulating the activation of RLC in skeletal muscle [79,80] and suggest that OVX may affect the activation of RLC in response to an acute exercise stimulus. It may be that the increased fat mass of the OVX group (i.e., lower muscle mass relative to body weight) contributed to the reduced maximal running capacity, an idea supported by the fact that the total workload during the maximal running test was not significantly different in the sham and OVX groups in the POST measurements despite the lower running distance achieved by the OVX group. Indeed, it has been observed in humans that excess body weight can influence distance run performance (Cureton et al., 1978 ).

### 4.4. OVX affected the acute response to exercise differently across brain regions

Regarding brain region specific findings, OVX was associated with lower serotonin metabolite 5-hydroxyindoleacetic acid (5-HIAA) level in brain NA, in accordance with earlier works [27,28]. As in the case of energy intake, the level of neurochemical markers following OVX may change in a time-dependent manner. For example, a recent study showed that in mice, OVX was associated with reduced hippocampal 5-HIAA at 1 week, but not 6 weeks after the surgeries [82]. However, another study in mice observed that 5-HIAA levels in HC remained reduced even after 28 weeks following OVX [83]. In our study (∼16 weeks post-OVX) we also observed a lower level of 5-HIAA following OVX, but this was true in the NA, and not in the HC brain region.

Contrary to previous literature, our results show that acute exercise had no effect on the level of serotonin in the sham group, while in OVX group it did affect brain serotonin level, albeit in a brain region-specific manner. Based on prior research, exercise, particularly acute bouts, increases the serotonin and 5-HIAA levels in the whole brain as well as the HC [for a review, please see [12]], while no data are available specific to the NA. Our study is the first to show that acute exercise is associated with higher serotonin levels in NA only in the OVX group. We speculate that this response observed in the OVX group may indicate a compensatory mechanism caused by the serotonin deficiency in NA associated with the loss of ovarian hormones. Noteworthily, exercise had the opposite effect on serotonin levels in HC for the OVX animals. Since serotonin is suggested to modulate fatigue upon prolonged exercise, the difference in serotonin response to exercise may contribute to the animals’ experienced exhaustion during maximal running tests [12].

Brain levels of serotonin, dopamine, and norepinephrine have all been linked to fatigue, thereby regulating physical performance [84]. Animal experiments indicate that increased serotonin reduces performance [85,86], while increased dopamine and norepinephrine increases performance [87,88]. Due to their interaction, serotonin/dopamine and possibly serotonin/norepinephrine ratio have been suggested to be more relevant for determining fatigue than analyzing only one neurochemical marker. However, we did not observe an effect of OVX, acute exercise or their interaction in serotonin/dopamine or serotonin/norepinephrine ratios in NA or HC. Also, when looking at the whole study population, we observed no significant correlation between the maximal running capacity results (POST, END) and serotonin/dopamine ratios in the two brain regions. However, consistent with previous observations, among the OVX group, there was a negative correlation between maximal running capacity (END) and serotonin/dopamine ratio in HC.

We noted a positive association between dopamine level in HC and maximal running capacity (POST) which has been shown previously [8]. In addition, HC dopamine level had a positive association with energy expenditure and markers of mitochondrial content (OXPHOS) in the retroperitoneal adipose tissue, which we believe to be the first report of a relationship between adipose tissue metabolism and HC levels of dopamine. Another relationship between brain neurotransmitter levels and body weight/adiposity was observed such that norepinephrine level in the NA was negatively associated with body mass and retroperitoneal adipose tissue mass. Given that brain-derived norepinephrine stimulates UCPs, these findings do provocatively suggest that OVX-mediated changes in brain neurochemical levels may contribute to changes in basal energy expenditure, a hypothesis that was not tested in the present study yet warrants further study.

### 4.5. Strengths and limitations

This study had many strengths but was not free of limitations that should be considered. The present study was designed to investigate the effects of OVX on maximal running capacity in adult female rats. In OVX literature, an extensive disagreement exists as to the ideal age at which rats should undergo the OVX surgery in order to mimic human menopause. Many studies consider animals adult once the animal is sexually mature, which does not correspond to the age when the animal is fully grown, relative to humans. Hence one key strength of the current study is that the animals were fully grown adults (∼7 months old) before they underwent the surgeries. The second strength of the study is the implemented allocation of the animals into the four study groups matched for body weight and maximal running capacity. Moreover, there were no differences observed in energy expenditure or spontaneous activity between the sham and OVX groups at baseline. A limitation of the current study is that the energy intake was not followed immediately after the OVX surgeries, where there may have been a transient increase in the energy intake of OVX rats. However, the current study’s focus was the long-term effects of OVX and the acute effect of exercise. It is also worth noting that OVX surgery causes a rapid decrease of systemic estrogens, and hence cannot be considered as an optimal model of human menopausal transition. Yet the long-term effects that OVX results in can represent some of the phenomena that takes place in women at the postmenopausal stage, such as increased fat mass and HDL-C levels, as reflected in our results.

### 4.6 Conclusions and future directions

In the adult female rats studied herein, OVX effectively mimicked menopause, as indicated by an increase in body mass during follow-up and higher fat mass, and HDL-C levels compared with the sham group. The changes in body mass and composition seemed to be driven by a decrease in basal energy expenditure. We report that OVX reduced maximal running capacity, but this was not accompanied by changes in muscle mass or pRLC in the gastrocnemius muscle. OVX was, however, associated with significantly lower brain serotonin metabolite 5-HIAA levels, confirming prior work by other groups. We extend those observations, demonstrating for the first time that the serotonin response to acute exercise differed in the two brain regions studied, HC and NA. These differences in the brain neurochemical response to an acute bout of exercise may be mechanistically related to the known OVX-associated changes in running behavior. These changes may potentially play a role in the reduced maximal running capacity, as well as the OVX-induced increases in body and fat mass. This may be due to the relationship between these neurochemical changes and whole-body energy expenditure.

## Acknowledgements

We would like to thank Mervi Matero, Suvi Kaura-aho, Eliisa Kiukkanen and the staff of the Animal unit at the University of Jyväskylä for the excellent care of the animals during the study. We acknowledge Timo Rantalainen for the mathematical equation for calculating work and Erik Niemi for his help with the data visualization. We also thank the laboratory staff at the Faculty of Sport and Health Sciences for their invaluable assistance in the data collection and analysis.

## Author contributions

SK, VV-P, and TAN conceptualized the study. EL, TAN, and SK performed the animal experiments. EL, TAN, LY-O, AJ, AL, and SK were in charge of the methodology and harvested the samples. EL, AJ, JEK and SK performed the formal analysis. EL, JEK and SK did the visualization of the results. SK was responsible for the funding, supervision, and project administration. EL, TAN and SK wrote the original draft and all authors reviewed and revised the manuscript before approving the final version.

