## Supplementary figures and images for "Estrogen deficiency reduces maximal running capacity and affects serotonin levels differently in the hippocampus and nucleus accumbens in response to acute exercise"

### Supplementary Figure 1

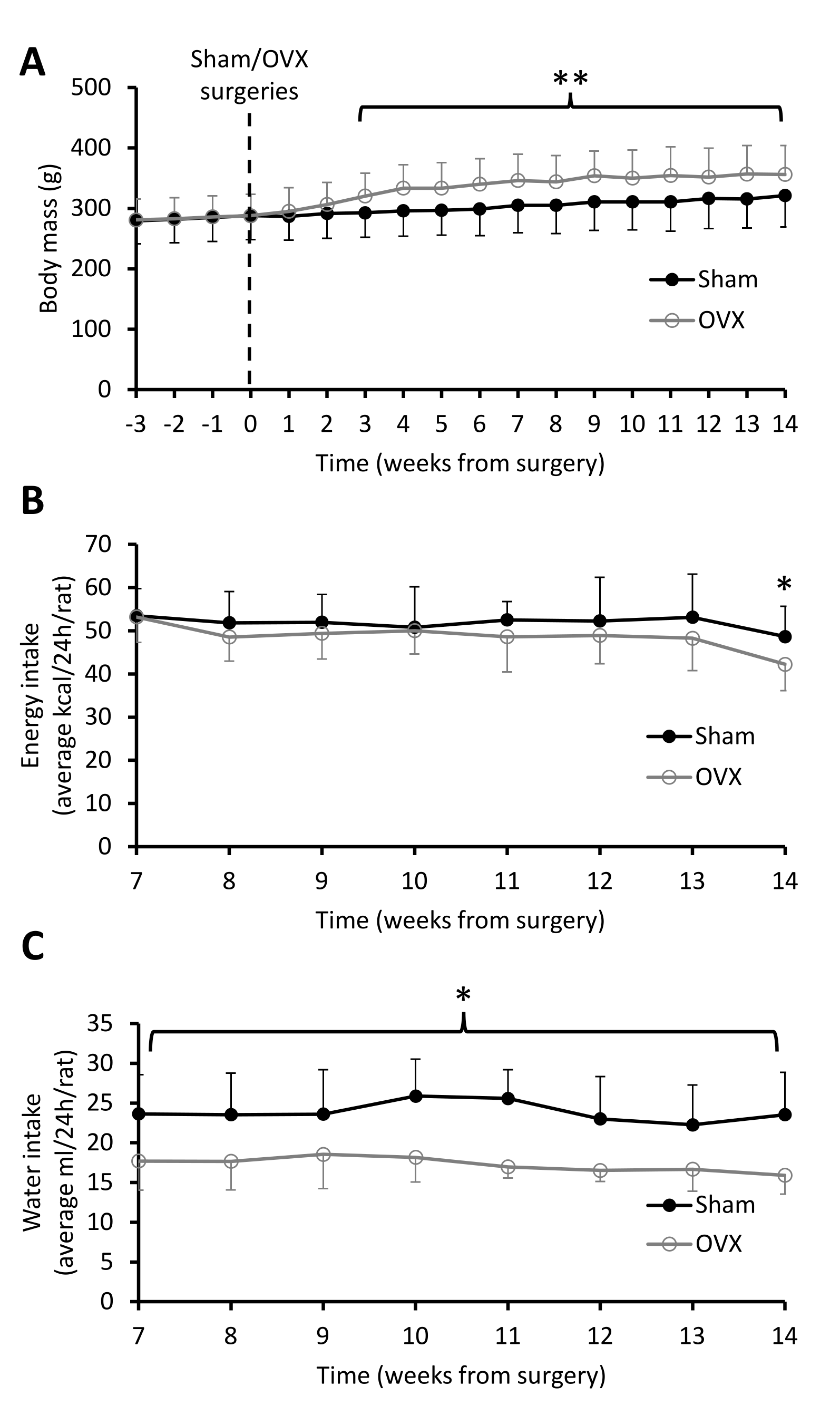

### Supplementary Figure 2

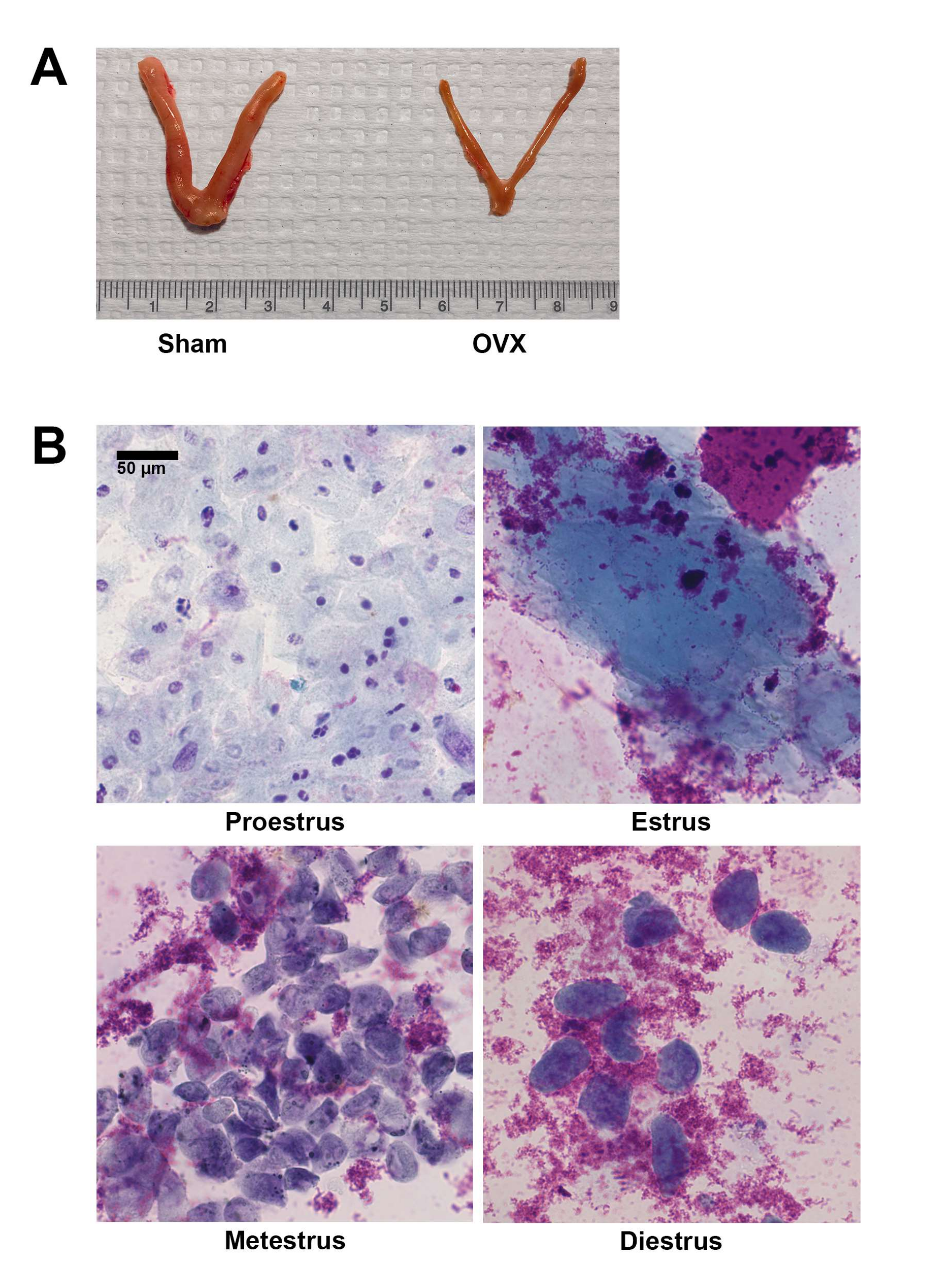

### Supplementary Figure 3

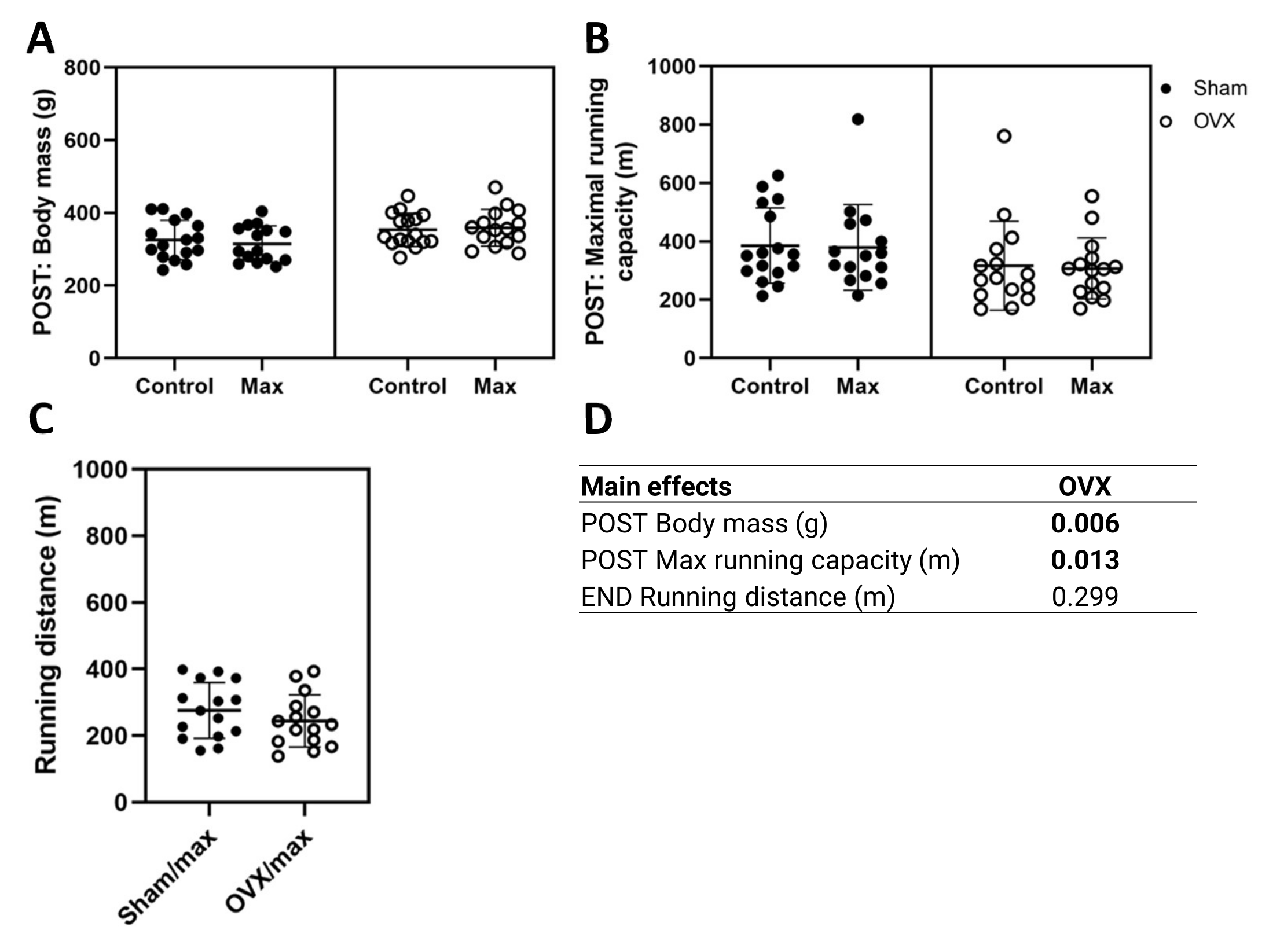

### Supplementary Figure 4

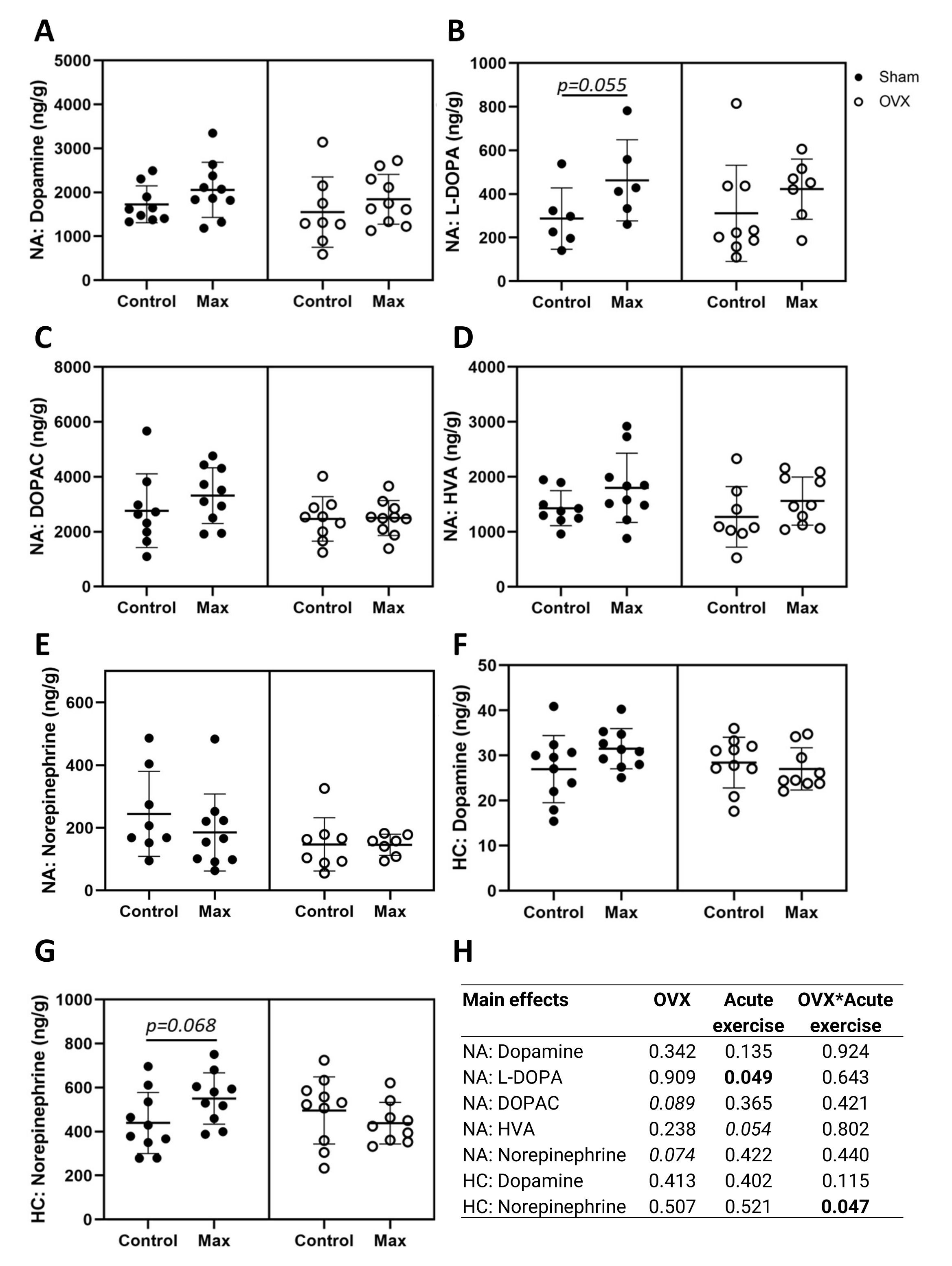

### Supplementary Figure 5

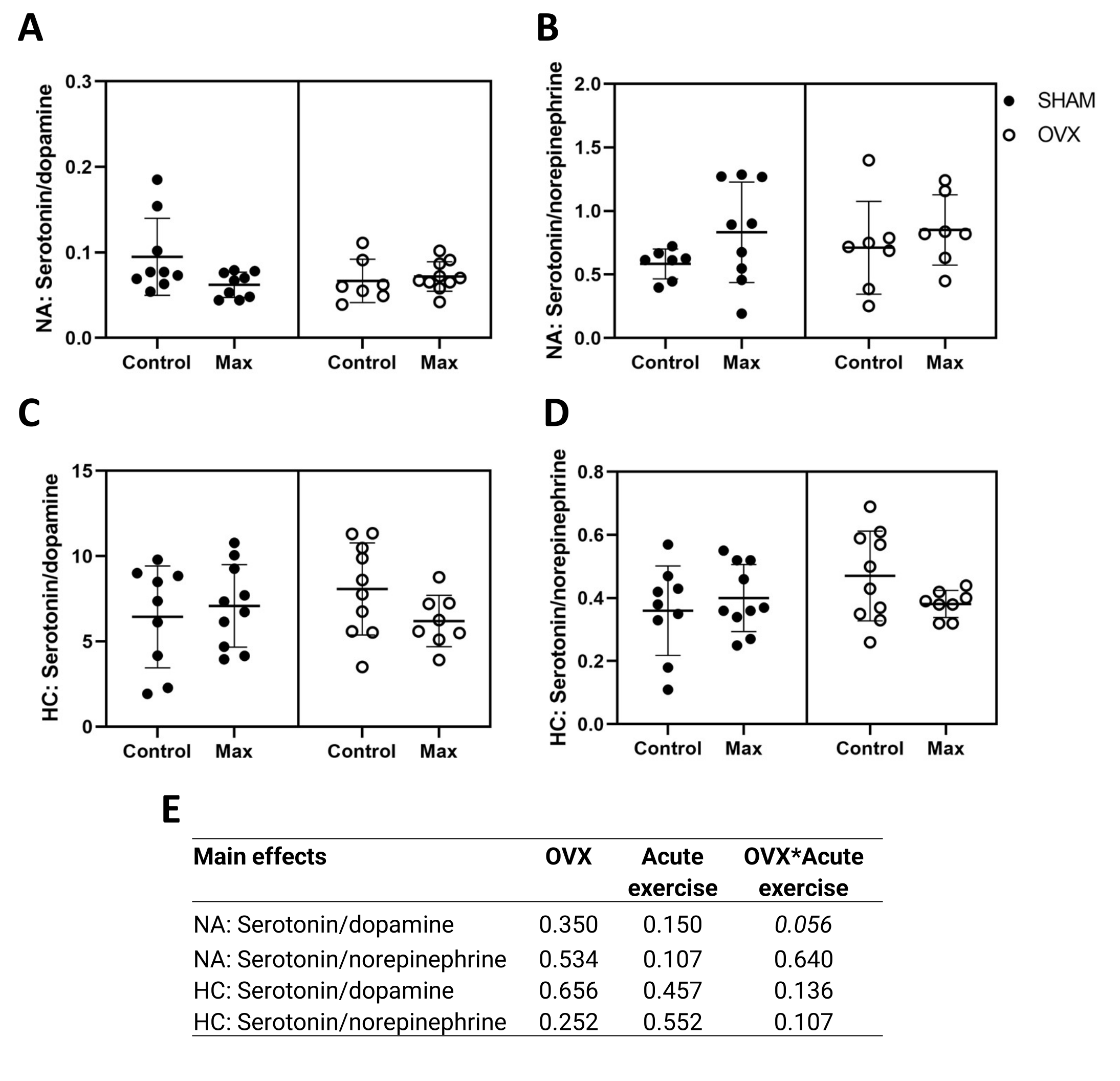

### Supplementary Figure 6

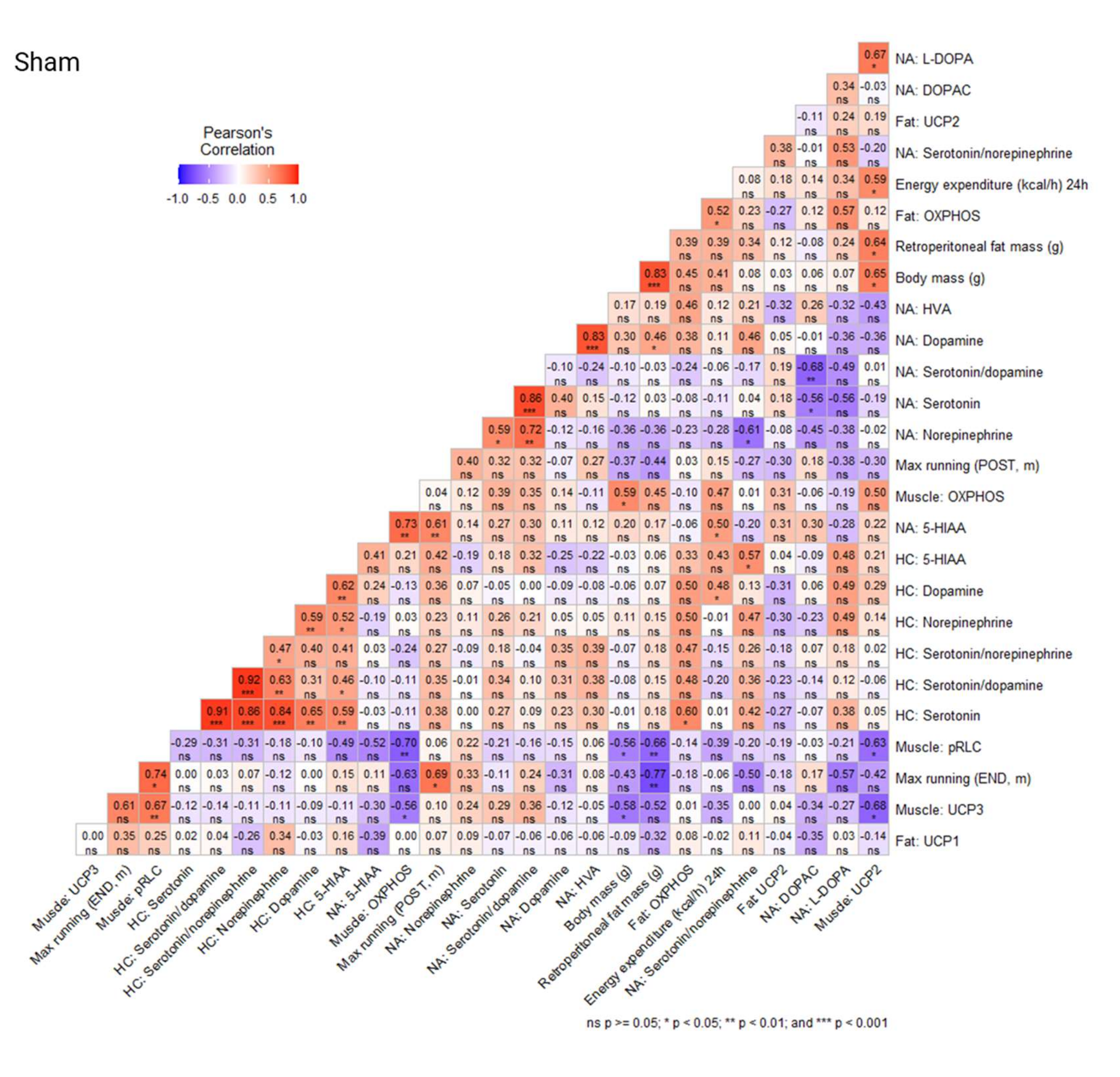

### Supplementary Figure 7

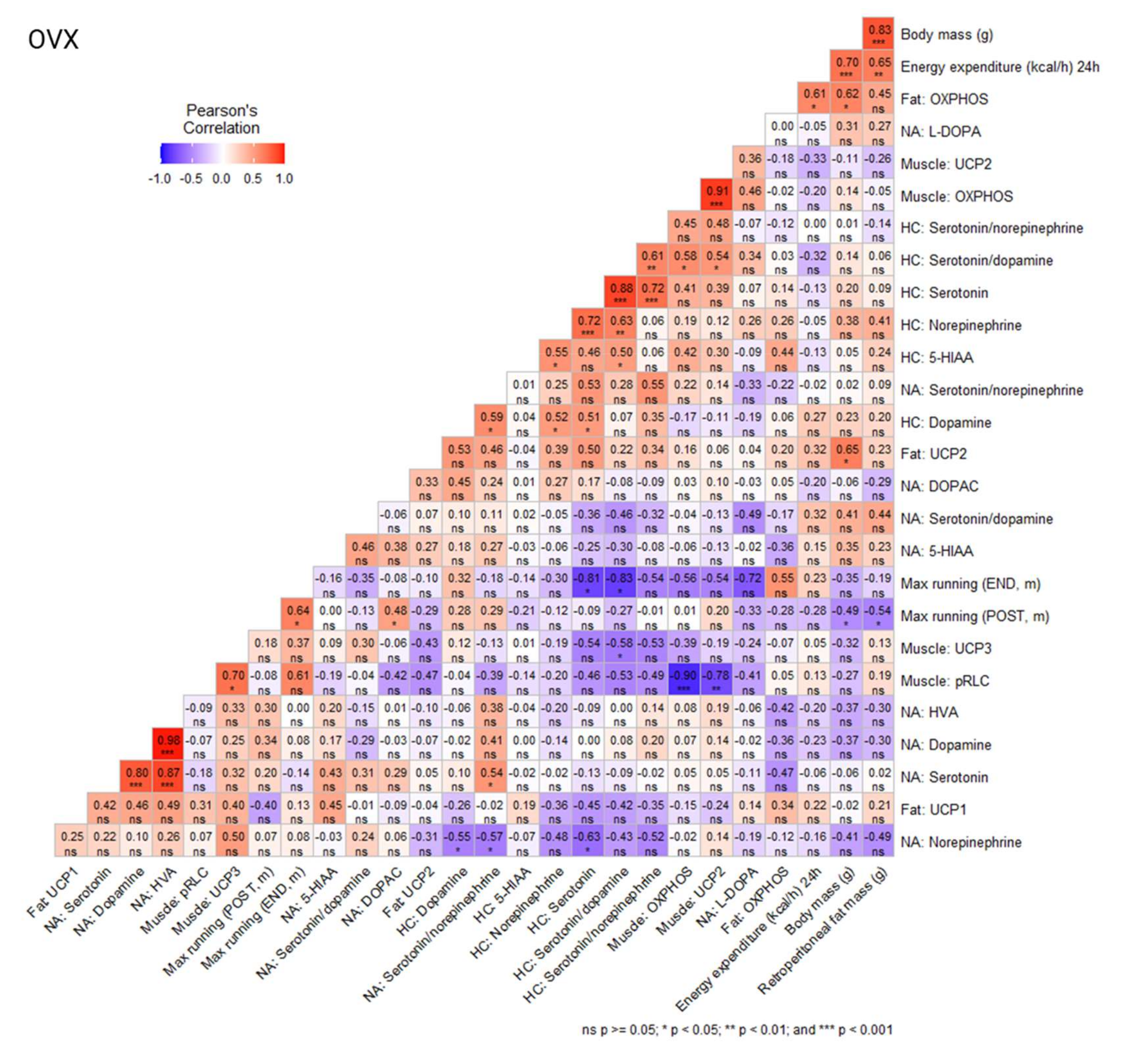

### Supplementary Figure 8

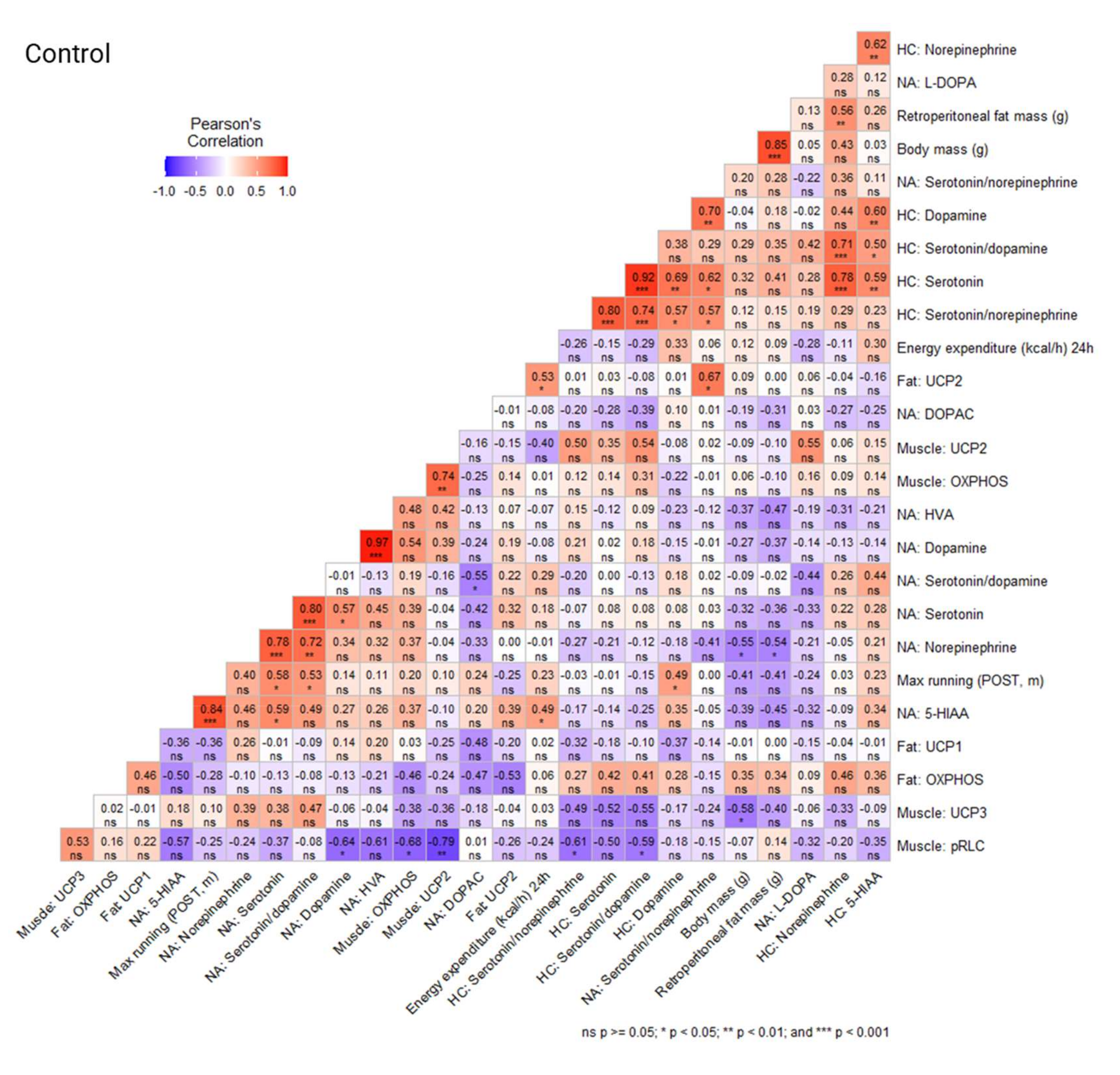

### Supplementary Figure 9

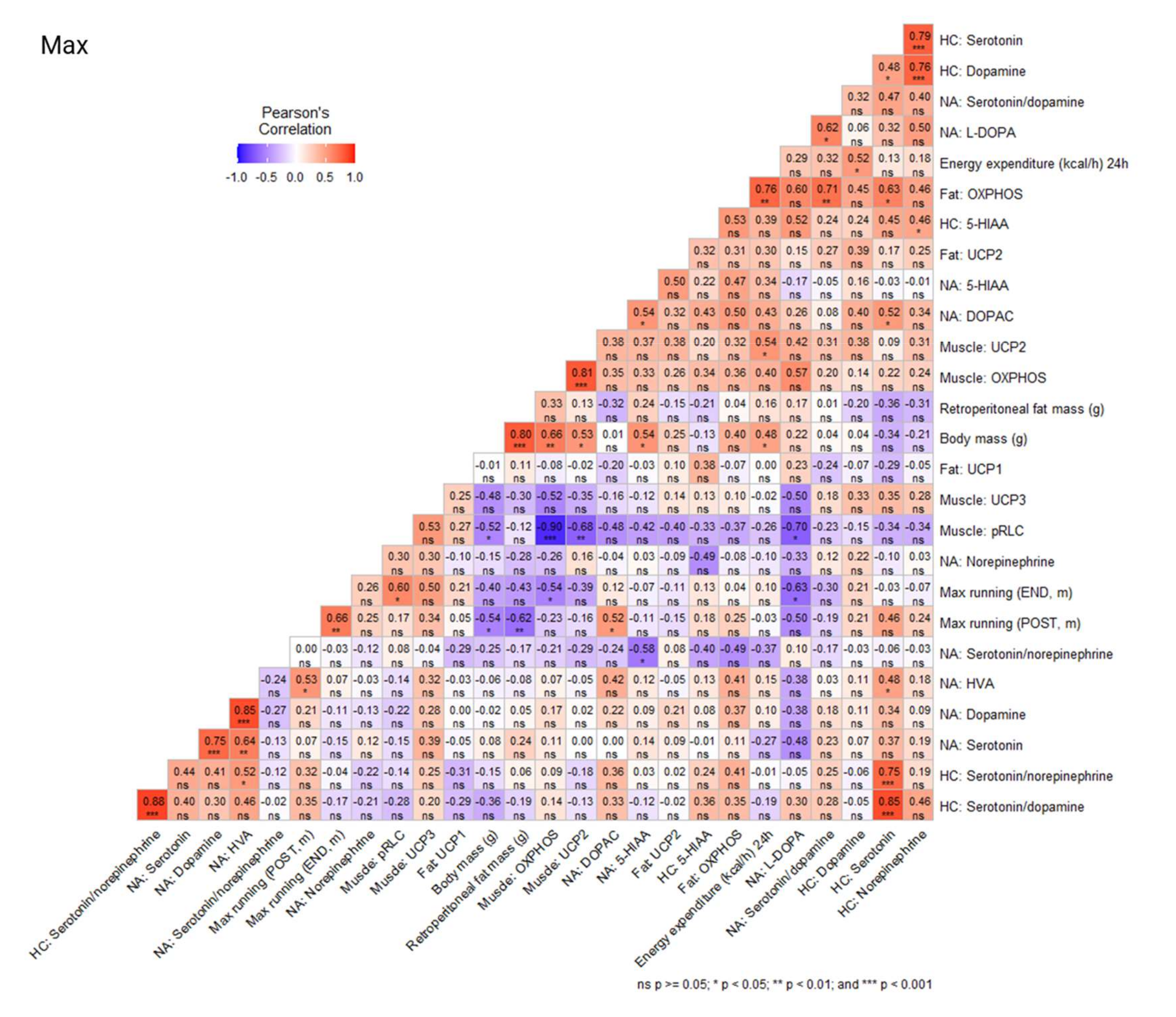
